# Seasonality, land use, and host diversity shape microbiome-pathogen interactions in wild populations of *Arabidopsis thaliana*

**DOI:** 10.1101/2025.06.29.662221

**Authors:** LP Henry, A Rat, E Laderman, R Lion, B Mayjonade, TEAM PATHOCOM, F Roux, D Weigel, J Bergelson

## Abstract

The microbiome often protects plants against pathogens, but most findings are limited to controlled experiments in the lab. In the context of wild populations, one key challenge is to understand sources of variation that impact the commensal microbiome, which in turn shapes the degree of protection. Here, we surveyed both disease symptoms and microbiomes from wild populations of *Arabidopsis thaliana* over four consecutive seasons (fall/spring) across three different land use types. Land use types varied in the extent of anthropogenic influences and included forest meadows, human-impacted fields adjacent to agriculture or municipal parks, and highly disturbed habitats near railroad tracks. By building an integrative map of abiotic and biotic variables, we find that a key predictor of disease was biodiversity across ecological scales. Plant communities with higher diversity were associated with reduced disease burden in *A. thaliana* populations but also increased diversity within the microbiome of *A. thaliana*. This increased microbial diversity was additionally associated with less disease in *A. thaliana*. However, the diversity-microbiome-disease relationships were all sensitive to season and further modulated by land use. Taken together, our work highlights the importance of anthropogenic change reshaping species interactions across ecological scales to impact disease risk in wild plant populations.

## INTRODUCTION

Rapid anthropogenic change is reshaping ecological processes at multiple scales that can impact infectious disease dynamics [1–3]. For example, warmer temperatures can exacerbate disease by increasing pathogen growth rates, while shifting plant physiological states that dysregulate the immune system and diminish defense responses [4]. In sum, anthropogenic change will alter many abiotic and biotic factors, and these disturbances are often implicated in reshaping the networks of ecological interactions [5]. Many pathogens are generalists and shared across species [6,7], and the impact of anthropogenic change on biodiversity and subsequent alteration of ecological networks may have potent impacts on disease dynamics.

On the other hand, the links between biodiversity and disease remain poorly understood. The dilution effect proposes that higher species diversity in communities dilutes disease risk by changes in community composition, context dependent variation in immune competence, and life history traits that decrease pathogen spread through a community [1,8,9]. Indeed, farmers have long recognized that intercropping and increased biodiversity in their fields is associated with lower disease incidence [10]. On the other hand, higher biodiversity is also often associated with higher pathogen richness and may be hotspots for pathogen emergence and spillover [11,12]. Nevertheless, an emerging synthesis suggests several key traits for understanding how land use change impacts disease dynamics. If land use changes increase the degree of habitat fragmentation, increase plant density, and decrease community diversity, then disease risk can be amplified [3]. However, the risk of disease spread will depend on factors that impede or facilitate microbial transmission within and between plant species [3,13,14].

Microbiomes have emerged as an important factor in protecting plants against pathogens [15–17], offering potentially new approaches for combating emerging pathogens [18,19]. The protection is thought to result from microbe-microbe resource competition or intermicrobial warfare, both of which inhibit pathogen colonization and growth [15,20]. Microbes can also indirectly modulate the host immune system and thus alter the effectiveness of defense against pathogens [21,22]. However, these findings are limited to controlled experiments in simplified ecological contexts. In realistic ecological contexts, plant disease dynamics are very different. Disease is often less severe in wild plants than in agricultural settings [23,24], and when disease outbreaks do occur, the incidence of coinfection is higher in wild populations [25]. While microbiomes are generally protective, interactions between the commensal microbiome and pathogen can instead lead to overall increases in pathogen diversity and burden [26,27]. Thus, microbiome-pathogen interactions potentially shape disease dynamics, but the impact remains poorly understood in naturalistic ecological contexts.

Understanding these complex microbiome-disease dynamics requires characterizing wild populations for both disease dynamics and microbiome variation. *Arabidopsis thaliana* is an excellent model as it inhabits a broad range of environments and co-occurs with diverse assemblages of other plant species. In the wild, *A. thaliana* harbor diverse microbiomes, shaped by a combination of host genetics [28,29], abiotic conditions [30], growing season [31], and the surrounding plant community [32,33]. In the wild, members of the *A. thaliana* microbiome can promote plant growth and can also protect its host against pathogens, such as *Pseudomonas* [32]. Beyond single pathogens, a humpbacked biodiversity-disease relationship between commensal microbes and pathogen diversity was observed In a survey of natural *A. thaliana* populations in France [26]. These complex dynamics suggest that a range of outcomes is possible, with commensal microbes either facilitating or suppressing pathogen diversity. Thus, it is clear that microbiome-pathogen interactions shape disease dynamics, but the impact remains poorly understood in naturalistic ecological contexts.

Here, we sampled wild *A. thaliana* populations in western Michigan across fall and spring for two years. *Arabidopsis thaliana* populations were sampled from different land use types that ranged in anthropogenic influence, from forests to plots immediately adjacent to railroad tracks. We characterized the abiotic and biotic environments to identify drivers of microbiome variation and associations with disease in the phyllosphere. If anthropogenic influence is restructuring ecological interactions, then we expect to see differences in microbial sharing between the surrounding plant community and focal *A. thaliana* stands across land use types. We then relate how these microbial dynamics are associated with variation in disease across the landscape.

Together, our results highlight how anthropogenic change shapes ecological interactions across biological scales, from within the microbiome to between plant species to overall disease dynamics.

## METHODS

Please refer to the supplement for full methodological details.

### Site characterization and sample collection

*Arabidopsis thaliana* populations were sampled from 19 sites in southwest Michigan in the fall and spring between Fall 2021 and Spring 2023 (Fig. 1A, Supp. Table M1 for site details, dates, and sample sizes); we note that Site 17 was only sampled once and subsequently replaced with Site 19. As *A. thaliana* at these sites are winter annuals, collections were timed to start shortly after germination in November and harvesting was completed in the spring, after bolting reproductive stalks but before flowering and senescence.

**Fig. 1:**
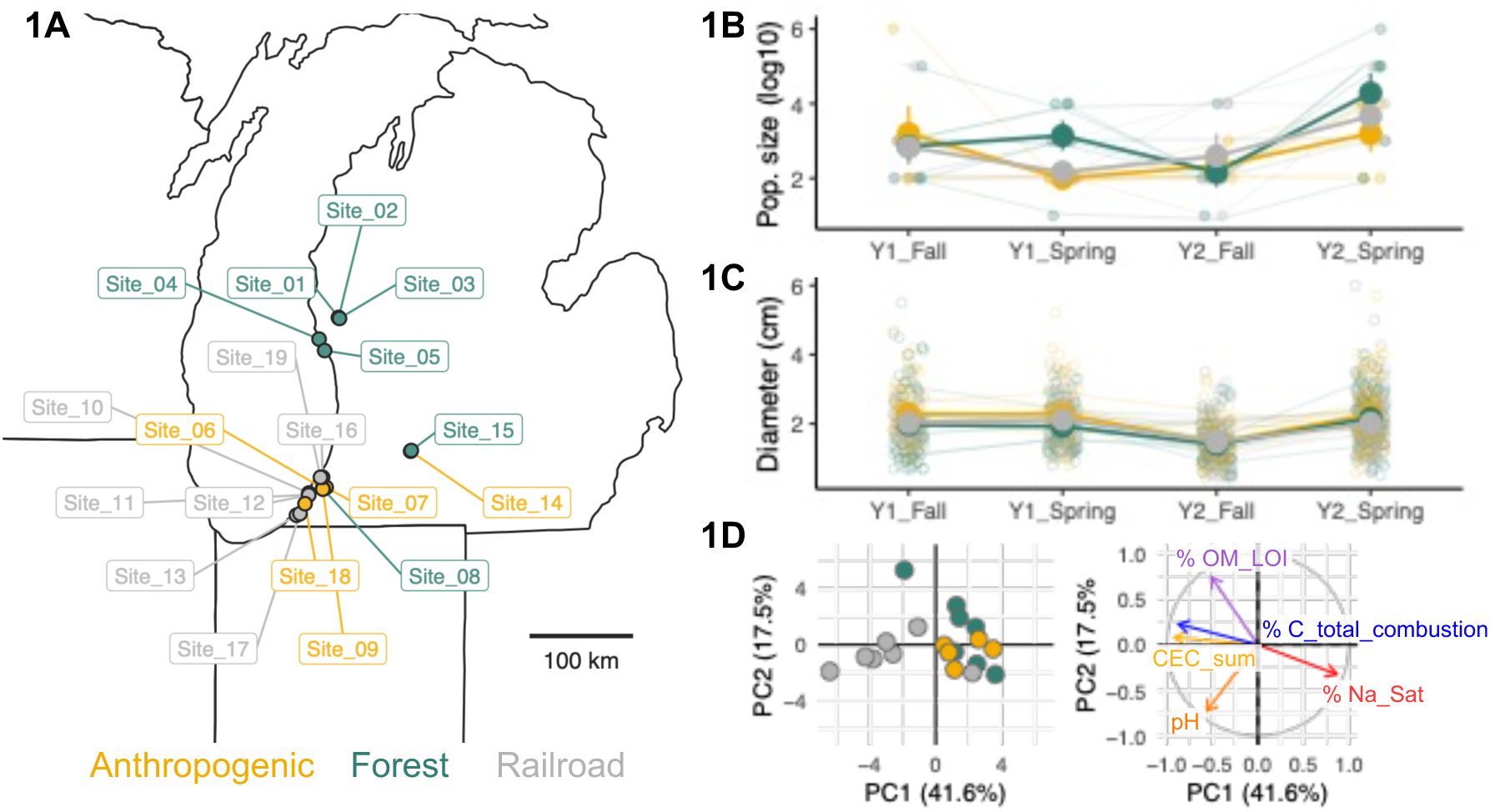
Survey of *A. thaliana* populations. A) Map of populations near Lake Michigan, USA, color-coded by land use. B) Population size across seasons, colored by land use type. C) Size of individual plants across seasons. For B and C, smaller hollow circles and thin lines represent each population, while solid larger circles, thick lines represent the average from each land use category, with standard error. D) PCA biplot showing soil chemistry results, with sites colored by land use on the right, and the left panel showing the top five most important loadings (text color matches arrow color to aid in visualization).

Sites were chosen to encompass the range of environments in which *A. thaliana* are commonly found in southwest Michigan, and the sites included populations previously identified for long-term genetic studies [34–36], including the 1001 Genomes Project [37]. For each site, two transects were established to capture the range of micro-environmental variation, and each subsequent sampling returned to approximately the same transect locations within the site.

Transects varied in size, ranging from 1.5 m to 17.5 m in length and with widths deviating as much as 1.5 m from the center of the transect. The length of transects was adjusted so that we could always collect 8-12 *A. thaliana* individuals. As we describe below, all plants were scored for disease symptoms, and approximately half of them were processed in the field to enrich for the endophytic microbiome. in an alternating patterns, plants were chosen to be enriched for endophytic microbiomes, to ensure that we did not bias microbiome samples in some way across potentially unknown environmental gradients. As we describe below, we further collected soil to assay for soil chemistry properties and microbiome content. We also characterized the companion plant community and their microbiomes. For all analyses, data from both transects were pooled per site and season, as our analyses focused on differences between sites.

We classified the land use type for sites into forest (N=7), anthropogenic (N=5), or railroad (N=7). Railroad sites were directly adjacent to active railroad tracks, while anthropogenic sites were often near agricultural fields or municipal parks. We used imperviousness data [38] to quantitatively distinguish forest and anthropogenic sites. Imperviousness data was determined at different spatial scales (every 50 m from 50-500 m, then 100 m up to 1 km). Forest sites were those with <5% imperviousness at all spatial scales, and the remaining non-railroad sites were classified as anthropogenic (Supp. Fig. M1); railroad sites had the highest imperviousness percentages at all spatial scales.

At each site, we characterized the abiotic environment relevant to plant-microbe interactions [33] using mean temperature on the day of sampling, cumulative precipitation in the three weeks preceding sampling, and the first two principal components of soil chemistry properties. Soil chemistry analysis was performed by A&L GreatLakes Laboratories. For the biotic environment, we focused on the companion plant community that co-occurs with *A. thaliana*. To further characterize companion plant communities at each sampling, two 50 cm x50 cm quadrats were placed in the approximate center of each transect, in areas with at least one *A. thaliana* individual within the quadrat. Percent coverage of the community was assessed using the quadrats. We then identified morphotypes as proxies for plant species, counted each morphotype, and saved a voucher for molecular identification using the chloroplast marker, *matK [39,40]*. Community diversity was pooled from the two transects, and analyses used the genus-level *matK* assignment. Bray-Curtis dissimilarity was used to partition variance in compositional changes due to land use type, and Shannon diversity was used as the primary alpha-diversity measure in subsequent analyses.

We collected the above-ground rosettes of 16-24 *A. thaliana* individuals at each site per sampling period (N=1,327 plants total). The distribution of *A. thaliana* population size was approximated at a log_10_ scale. All plants were visually scored for the following disease symptoms [41]: chlorosis, necrosis, hypersensitive response (HR)-like lesions, *Albugo* presence, *Hyaloperonospora* presence, general fungal damage, and herbivory. A composite disease score (excluding herbivory) was built using the presence/absence of each symptom using principal components analysis. Other plant phenotypes measured were diameter, number of leaves, and bolting.

The leaf microbiome is composed of both epiphytic microbes that live on the surface along with endophytic microbes inside the leaves [42]. Most pathogens are endophytic, but the endophytic compartment also contains many commensals and thus likely captures relevant pathogen-commensal interactions. Approximately half of the *A. thaliana* samples were randomly selected to be enriched for the endophytic portion of the microbiome in the field by depleting the loosely-associated epiphytic microbes through a series of washes following established protocols [43,44]. Plants were briefly washed with 1X TE + 0.01% Triton-X, 80% ethanol, 2% bleach, and sterile water. Rosettes were blotted dry and placed immediately on dry ice until returned to the lab. We also collected a small portion of each non-*A. thaliana* morphotype to characterize the companion plant microbiome from pooled samples, as well as soil samples for microbiome characterization. Thus, microbiome analyses were focused on the three sample types: individual *A. thaliana* endophyte samples, companion plant pool, and soil.

### Microbiome characterization

We characterized the *A. thaliana* microbiome using host-associated microbe PCR (hamPCR) [45] to simultaneously assess microbial load and community diversity (see supplement for full details). hamPCR co-amplifies a microbial marker gene with a single copy host gene in a single PCR reaction and then incorporates Illumina adapters for high-throughput sequencing. We used hamPCR for both bacteria (*16S rDNA* V5V6V7) and fungi (*ITS1-2*) with *GIGANTEA* as the host gene; the same amplicons were used for soil and companion plants without *GIGANTEA*. Sequencing was performed on the Illumina NovaSeq platform, 1x300 bp at the NYU Genomics Core.

Microbial load is calculated by dividing the microbial reads by the *GIGANTEA* reads, and composition is determined using tools implemented in QIIME2 v2023.2 [46] and phyloseq [47]. ASVs were called using DADA2 [48] and taxonomically assigned using classify-sklearn Bayes classifier [49]. 16S rDNA were classified using the Greengenes database [50] and ITS sequences were classified using the Unite database [51]. Fungal trophic modes were assigned using the FunGuild package [52]. Microbial load and composition were treated as separate traits, and we did not scale endophytic microbial composition by load. By treating microbiome composition as a distinct trait, we could compare relative abundances of particular microbes in the endophyte samples and in the sample types without measured microbial load (companion plants and soil).

For the *A. thaliana* endophyte samples only, we assessed finer-scale phylogenetic resolution by developing primers that target strain diversity within *Sphingomonas* and other closely related genera (*mmdA* gene) and within the *Pseudomonas* genus (*HSD* gene). We chose to focus on these groups because *Sphingomonas* was the most abundant bacterial taxon and *Pseudomonas* contains both plant growth promoting strains [15] and important pathogens of *A. thaliana* as well as many other plants. *mmdA* and *HSD* amplicon libraries were prepared without co-amplifying *GIGANTEA*, and sequencing data were processed as stated previously for 16S rDNA and ITS, but taxonomy was assigned using a custom database. Sequencing was performed on the Illumina MiSeq platform, 1x300 bp at the NYU Genomics Core.

### Statistical approach

Given the complexity of the following analyses, we briefly outline our general approach here, but provide more specific tests in the results section (please see supplement for full details). We analyzed our data using mixed linear models and PERMANOVAs to identify associations between *A. thaliana* phenotypes (i.e., disease symptoms, microbiome variation) with environmental factors (see supplement for full details). For all mixed linear models, population was nested within the sampling year as a random effect. For PERMANOVAs assessing compositional differences (companion plant community and microbiome beta-diversity), population membership was used as blocks and year as a main term. For all microbiome beta-diversity analyses, we used a modified Aitchison distance [53]. While this distance metric is largely insensitive to read depth, we included read depth as a covariate, as rarefaction was not performed before ordination. Alpha-diversity metrics for all amplicons were rarefied to 1,000 reads for endophyte samples and 16,963 reads for companion plant and soil microbiomes (Supp. Fig. M2, supplemental methods for more detail). For models that focused only on endophytes (e.g., disease dynamics, microbial load and diversity), plant diameter was included as a fixed effect to control for differences in host size. As land use type and season often co-varied with many other factors, we primarily used these qualitative factors, but delineated when we used quantitative, site-level factors (e.g., companion plant community diversity, soil chemistry PC1, etc.). We often tested for interactions between season and land use, but if interactions were not significant, the interaction terms were dropped. Mixed effects models were implemented in lme4 [54]. Residuals were assessed for dispersion, normality, and outliers using DHARMa [55]. Beta-diversity analyses used the adonis2 implementation of PERMANOVA in vegan [56].

To link all the datasets and explore direct and indirect effects on disease dynamics, we used structural equation modeling to examine the impacts of land use, season, companion plant, and microbial variation (i.e., diversity and load). Path analysis was informed by results from mixed linear effects modeling, which suggested the importance of bacteria in shaping disease PC1 and fungi in shaping disease PC2. Given the strong effects of season and to better visualize patterns, we further separated path analysis between seasons. We constructed a model that included direct effects of non-microbial variation (land use type, season, population, year, plant size, companion plant Shannon diversity) and microbial variation (bacteria/fungal Shannon diversity, bacterial/fungal load) on disease. Then, we modeled the indirect effects on non-microbial variation on microbial variation. Structural equation modeling was performed using Lavaan [57].

## RESULTS

### Land use shapes abiotic and biotic factors in A. thaliana populations

Across the 19 populations sampled, *A. thaliana* population sizes varied idiosyncratically across years and season, with no significant difference between land use types (Fig. 1B, Supp. Table R1). *Arabidopsis thaliana* plant size increased in the spring, again with no significant differences between land use type (Fig. 1C, Supp. Table R2).

We then characterized how abiotic and biotic factors vary across land use types for *A. thaliana* populations. For abiotic factors, we focused on temperature, precipitation, and soil chemistry. Neither temperature nor precipitation significantly differed between the land use types (Supp. Fig. R1, Supp. Table R3). Soil chemistry differed between land use types, where railroad soils were separated from anthropogenic and forest soils along PC1 (Fig. 1D; full details in Supp. Fig. R2); railroad soils tended to have higher cation exchange capacity and organic matter levels.

For biotic factors, we characterized the co-occurring companion plant community, which potentially shares microbes with the focal *A. thaliana* individuals. The composition of companion plant communities was primarily structured by land use type (R^2^ = 0.09, F_2,66_ = 2.71, p=0.002), with soil chemistry, temperature and year explaining marginal, but significant amounts of variance (Supp. Fig. R3, Supp. Table R5). The most abundant plants were *Arenaria* sp., *Cerastium* sp., and *Veronica* sp., as well as genera that were unique to each land use type (Supp. Fig. R3). Alpha-diversity of the companion plant community did not significantly differ across land use types or seasons (Supp. Fig. R4, Supp. Table R6). Taken together, the different land use categories are associated with different abiotic and biotic factors.

### Land use modulates seasonal patterns of disease dynamics

We scored ∼1,300 *A. thaliana* individuals for disease symptoms across the four sampling periods. The most frequently observed symptom was chlorosis, followed by HR-like lesions and necrosis (Fig. 2A). Many *A. thaliana* individuals often featured multiple disease symptoms, with the most common co-occurrence being chlorosis, HR-like, and necrosis. Notably, we detected a strong seasonal association for plants with multiple symptoms, with most plants in the spring exhibiting at least two different symptoms (Fig. 2B). More so, the dominant symptoms exhibited different temporal fluctuations in the different land use types (Supp. Fig. R5). For example, the presence of HR-like lesions decreased in the spring for forest populations, but increased for railroad populations; anthropogenic sites did not display seasonal dynamics for this symptom.

**Fig. 2:**
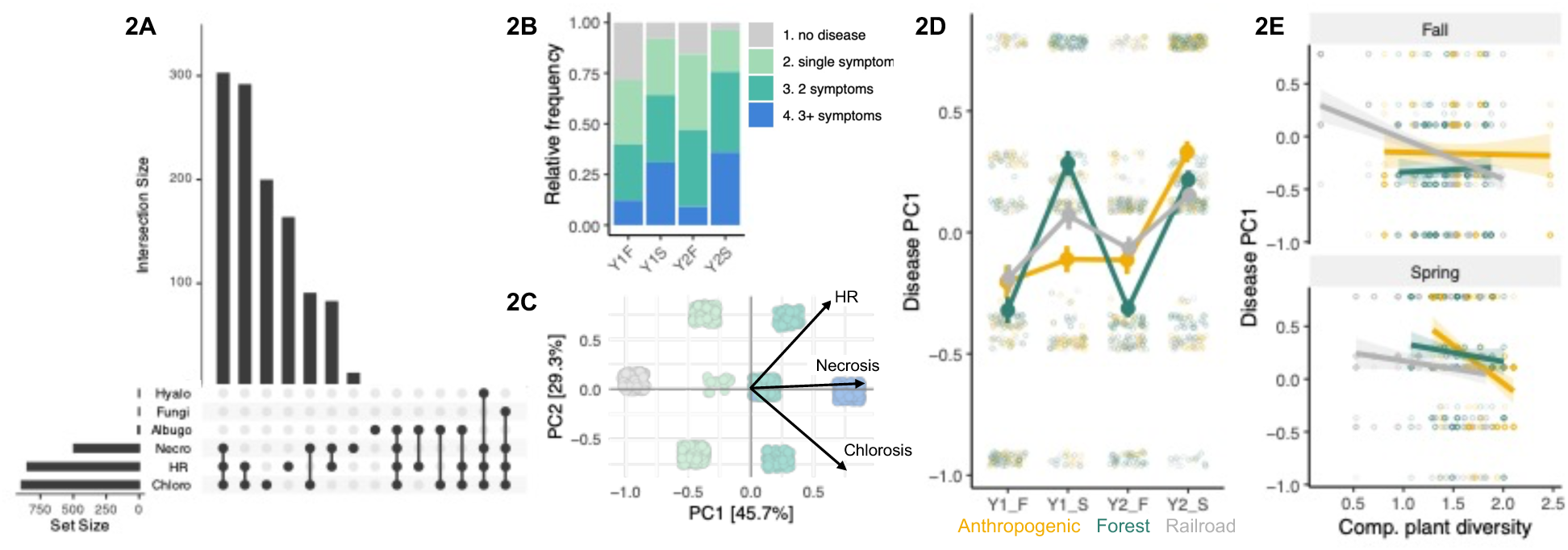
Interactions between land use and season shape disease dynamics. A) Upset plot showing the overlap between different disease symptoms and counts across all plants sampled. B) Relative frequency of plants with the number of symptoms observed. C) PCA showing composite disease score, where each point is an individual sample, colored by the number of symptoms observed that separate PC1. The arrows indicate the driving symptoms that separate PC2. D) Temporal dynamics in disease dynamics as indicated by the first principal component of the composite disease score. Smaller hollow circles represent individual plants, while the larger solid circles and thick lines represent the average from each land use category per season, with standard error. E) Companion plant community diversity is negatively correlated with disease PC1, but depends on land use and season. Small points represent individual plants, colored by land use.

To understand how land use influences disease symptoms, we first integrated the different disease symptoms into a composite disease score using PCA loadings. PC1 separated the single- and multiple-symptom plants, and PC2 separated plants by symptom types (Fig. 2C). With this composite score, we detected a significant three-way interaction between disease PC1 and land use, season, and Shannon diversity of the companion plant community (interaction Wald X^2^ = 33.3, p < 0.001, Fig. 2D-E, Supp. Table R7). Forest populations exhibited increases in symptoms in the spring, and these swings were greater than in railroad populations (Fig. 2D). Populations at sites with strong anthropogenic influence, however, did not display a seasonal pattern. Companion plant community diversity was negatively correlated with disease PC1 across all land use types in the spring but only in the railroad populations during the fall (Fig. 2E). In the spring, the relationship between disease PC1 and companion plant diversity was steepest for anthropogenic sites. Overall, we find that the diversity of companion plants impacted disease symptoms in wild *A. thaliana,* but this impact was sensitive to land use and season.

### Season strongly structures the endophytic microbiome of A. thaliana, but land use shapes microbial sharing with companion plants

We next assessed factors shaping microbial dynamics in *A. thaliana*. We explored three axes of variation in the bacterial and fungal microbiomes: 1) microbial load in *A. thaliana* endophyte samples (standardized microbial read number to host read number), 2) the drivers of beta- and alpha-diversity differences between and within sample types, and 3) microbial sharing between *A. thaliana*, companion plants, and soil.

Microbial load of *A. thaliana* endophytes was significantly higher in the spring than in the fall for both bacteria and fungi (bacteria: β = 0.89 ± 0.04 SE, p < 0.0001, fungi: β = 0.16 ± 0.07 SE, p = 0.02, Fig. 3A-B, Supp. Table R8); more variance was explained for bacterial load (marginal R^2^ = 0.29) than fungal load (marginal R^2^ = 0.01). Bacterial and fungal loads were positively correlated (β = 0.26 ± 0.03 SE, p< 0.001), although the strength of this correlation differed between fall and spring (interaction Wald X^2^ = 23.8971, p < 0.0001, Supp. Fig. R6, Supp. Table R9). Land use did not significantly affect bacterial or fungal load.

**Fig. 3:**
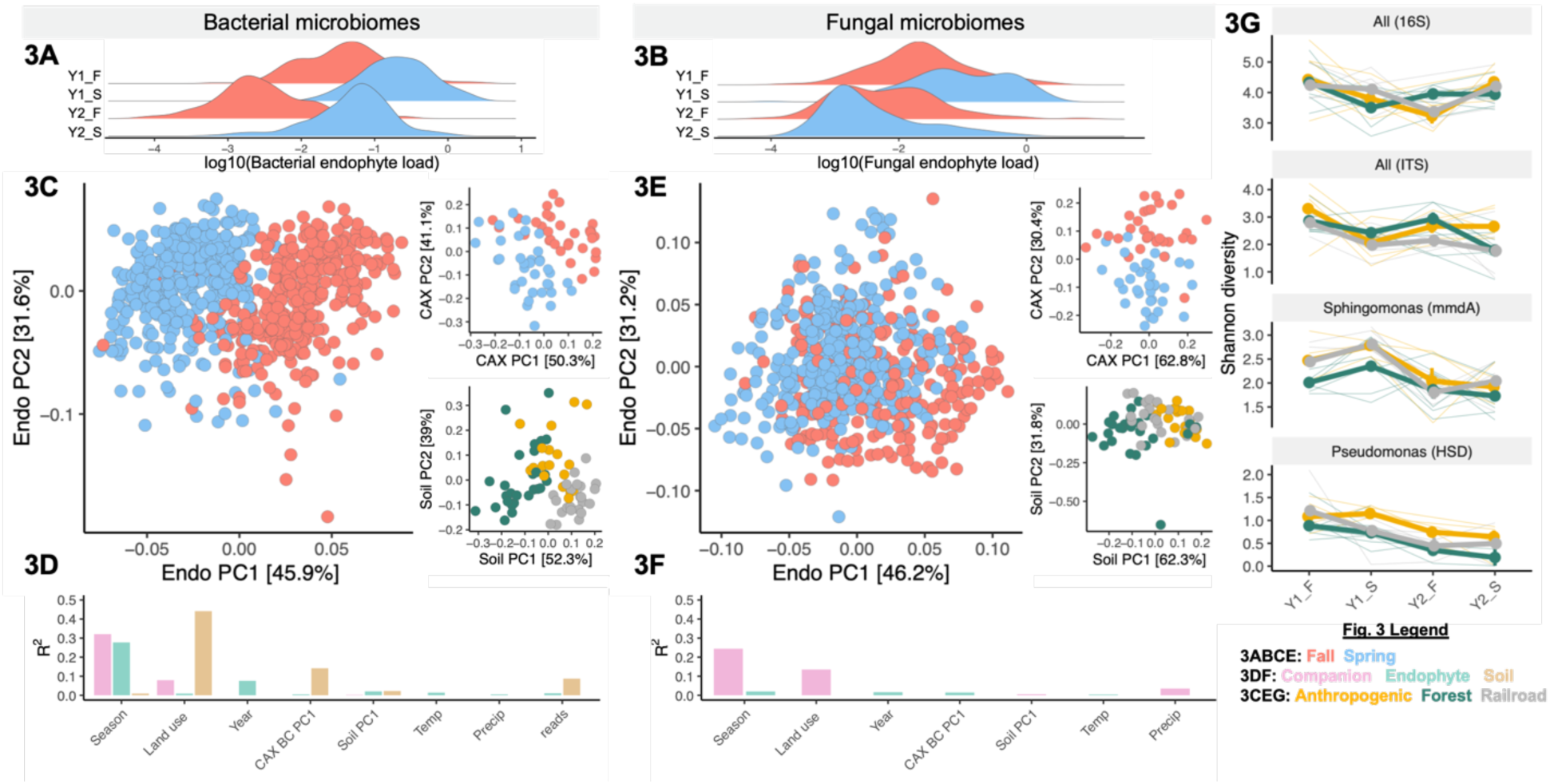
Season and land use impact microbiomes across sample types. A) Bacterial load in spring (blue) and fall (red). B) Fungal load in spring and fall. C) Ordinations for bacterial microbiomes in *A. thaliana* endophyte samples, companion plants (abbreviated as “CAX”), and soil based on modified Aitchison distance matrix. Points are colored by the variable that explained the most variance. D) Bar plots of the R^2^ value of each factor in PERMANOVA analysis for bacterial microbiomes, with color representing sample type (CAX BC PC1 = plant community beta-diversity using Bray-Curtis PCoA Axis 1). E) Ordinations for fungal microbiomes, colored by variables that explained the most variance. F) Bar plots of the R^2^ value of each factor in PERMANOVA analysis for fungal microbiomes, with color representing sample type. No significant variance as explained by any factor for soil microbiomes. 3G) Shannon diversity of each amplicon across the four sampling seasons, colored by land use. Thin lines represent each population, while thick lines and solid circles show mean, with standard error.

Next, we focused on drivers of compositional differences between sample types. Most microbes were found across all sample types, but their relative abundance differed (Supp. Fig. R7 for bacteria, Supp. Fig. R8 for fungi); these microbes are also commonly observed in wild *A. thaliana* populations [28,30–32]. Sphingomonadaceae were the most abundant bacteria across all sample types, while the most abundant fungal taxa varied (Table 1). Sample type only explained 8.1% of variance in bacterial microbiomes using a modified Aitchison distance metric (F_2,879_ = 50.7, p = 0.001, Supp. Fig. R9, Supp. Table R10). Less variance was explained for fungal microbiomes, though sample type explained the most variance (R^2^ = 0.04, F_2,760_ = 14.8, p = 0.001, Supp. Fig. R9, Supp. Table R10). We note that other beta-diversity metrics explained less variance, but show qualitatively similar results (Supp. Fig R10, Supp. Table R11). Alpha-diversity was highest in soil and lowest in endophyte samples for both bacterial and fungal microbiomes (Supp. Fig. R11, Supp. Table R12).

**Table 1:** Top three most abundant microbes for each sample type.

| ***Bacteria*** |  |
| --- | --- |
| **Endophytes** | Sphingomonadaceae (33.7%), Oxalobacteraceae (15.9%), Comamonadaceae (8.5%) |
| **Companions** | Sphingomonadaceae (25.4%), Oxalobacteraceae (10.3%), Comamonadaceae (8.7%) |
| **Soil** | Sphingomonadaceae (13.4%), Actinobacteria (6.3%), Beijerinckiaceae (5.6%) |
| ***Fungi*** |  |
| **Endophytes** | Mycosphaerellaceae (15.2%), Amphisphaeriaceae (13.5%), Helotiaceae (9.1%) |
| **Companions** | Didymellaceae (13.0%), Mycosphaerellaceae (10.5%), Cladosporiaceae (7.2%) |
| **Soil** | Nectriaceae (10.6%), Trichomeriaceae (5.3%), Helotiaceae |

While sample types generally shared the same microbes, different forces structured the microbiomes within each sample type. In these analyses, we included qualitative site descriptors (e.g., land use, season) as well as quantitative measures (e.g., companion plant community PC1, temperature, etc). Season structured beta-diversity in the endophytic bacterial microbiomes (R^2^ = 0.28, F_1,743_ = 349.3, p = 0.001) and companion plant microbiome (R^2^ = 0.32, F_1,67_ = 36.5, p = 0.001), while soil bacterial microbiomes were more strongly structured by land use (R^2^ = 0.44, F_2,67_ = 47.8, p = 0.001, Fig. 3C-D, Supp. Table R13). For fungal endophytes, season explained most of the little variation that could be explained (R^2^ = 0.02, F_1,624_ = 13.3, p = 0.001, Fig. 3E-F, Supp. Table R14). Like for bacteria, season structured the fungal microbiomes in companion plants (R^2^ = 0.32, F_1,67_ = 26.7, p = 0.001, Fig. 3E-F, Supp. Table R14). Other variables generally did not explain substantial variance structuring either bacterial or fungal microbiomes, with the exception that companion plant community PC1 explained significant variance in the soil bacterial community (R^2^ = 0.14, F_1,67_ = 30.7, p = 0.001, Fig. 3D-F, Supp. Table R13).

For alpha-diversity, we focused on the endophytes and expanded our dataset to include two additional amplicons, *mmdA* and *HSD*, which capture fine-scale diversity within *Sphingomonas* and *Pseudomonas*, respectively, allowing us to compare the response of microbiomes to land use at different phylogenetic scales. To test this, we compared patterns in temporal change for Shannon diversity for the different amplicons and found that a three-way interaction between amplicon, land use and season shapes temporal dynamics (three-way interaction Wald X^2^ = 15.4, p = 0.02, Fig. 3G, Supp. Table R15). At the community level (i.e., 16S and ITS), we saw few trends within land use types, except for consistent decreases in alpha diversity within forests in the spring. At the strain level, we found that *Pseudomonas* alpha diversity tended to decrease over sampling periods for all land use types. The pattern was less clear for *Sphingomonas*, in which alpha diversity increased in the Spring across all land use types in the first, but not second, year of our study. There were no effects of land use or season on alpha diversity of the fungal or bacterial microbiomes in the soil or companion plants (Supp. Fig. R12, Supp. Table R16); recall that we did not investigate *Sphingomonas* or *Pseudomonas* diversity in these sample types. Finally, we examined correlations between load and Shannon diversity to understand how land use affected these two aspects of microbial variation. Shannon diversity and load were weakly correlated for bacterial microbiomes, with the direction of the correlation differing between land use types (interaction Wald X^2^ = 13.0, p = 0.001, Supp. Fig. R13, Supp. Table R17). For fungal microbiomes, a strong negative correlation was observed between Shannon diversity and load, but without an effect of land use (β = -1.04 ± 0.06 SE, p < 0.0001, Supp. Fig. R13, Supp. Table R17). These high-load, low-diversity fungal microbiomes are not dominated by necrotrophs nor are the fungal taxa in these samples unique; the same fungal taxa are also observed across samples with higher diversity and lower load (Supp. Fig. R14).

Taken together, we conclude that season and land use structure the alpha-diversity of endophytic communities within *A. thaliana* endophyte samples, with differing impacts at microbial species versus strain levels, but no obvious effect for the soil and companion plant microbiomes.

The phyllosphere is assembled through microbial dispersal from the air, soil, and other co-occurring plants [42]. If land use is impacting the potential for microbial spillover from other sources, then we would expect to see associations between microbes in *A. thaliana* and microbes in companion plants or soil. We first examined this at the macro-community level to understand how the companion plant community affected *A. thaliana* microbial diversity. Indeed, companion plant community diversity, and thus the potential diversity of microbial sources, and *A. thaliana* endophyte bacterial and fungal diversity were positively correlated, but the correlation differed between land use types (bacterial interaction Wald X^2^ = 8.55, p = 0.01; fungal interaction Wald X^2^ = 6.49, p = 0.04; Supp. Table R18); there was no interaction with season. For bacteria, a positive correlation was observed for all land use types, but was steeper for anthropogenic sites (Fig. 4A). For fungi, the positive correlation was only observed in forest populations (Fig. 4B). Furthermore, *A. thaliana* microbial Shannon diversity was correlated with both microbial diversity from companion plants and soil, but the effects of land use and season differed between bacteria and fungi (Supp. Fig. R15, Supp. Table R19).

**Fig. 4:**
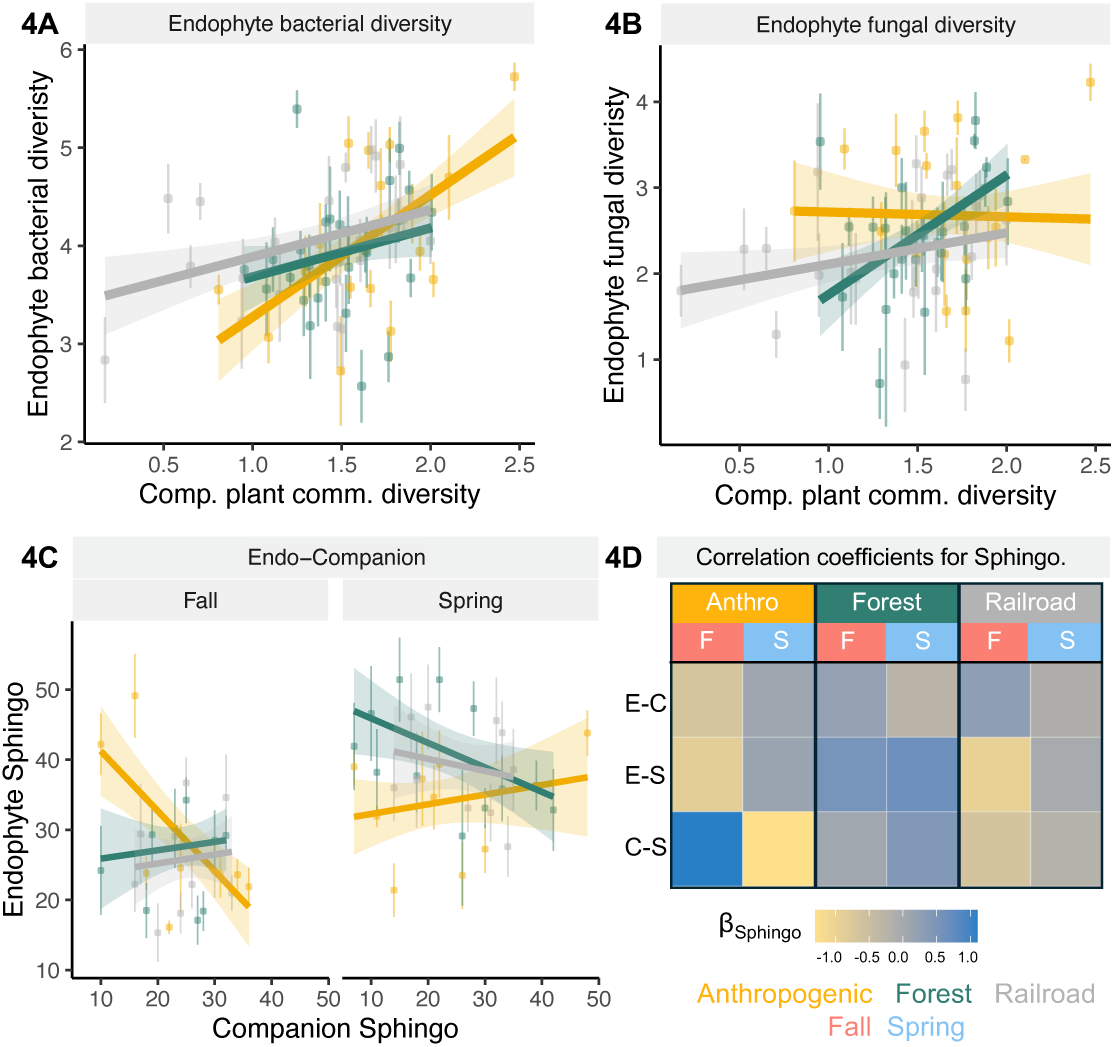
Land use impacts microbial diversity and sharing between sample types. For A-C, points represent the correlation between companion plant and mean (+/- standard error) for relative abundance in the endophyte samples for each population, colored by land use. A) *Arabidopsis thaliana* endophyte microbial diversity is positively correlated with the companion plant community diversity, but the slope differs between land use types. B) *Arabidopsis thaliana* fungal diversity is also positively correlated with companion plant community diversity, but not for anthropogenic sites. C) Correlations for the relative abundance of the dominant bacterial family, Sphingomonadaceae, between companion plant and *A. thaliana* endophytic microbiomes. Other sample types are in Supp. Fig. R17. D) Boxes represent the Sphingomonadaceae correlation coefficients (β) for a given sample type pair. Y-axis titles are abbreviated comparisons for E-C: endophyte-companion, E-S: endophyte-soil, and C-S: companion-soil.

Next, we assessed microbial sharing between sample types using correlations for the dominant bacterial taxa in endophytes, Sphingomonadaceae, across land use types and seasons. We compared *A. thaliana* microbes with those in companion plants because of the detected effect of companion plant community diversity on microbial diversity (Fig. 4A), and also with soil microbes because of the strong effect of land use (Fig. 3D). We did not include fungi because the dominant taxa varied across sample types more than in bacterial microbiomes, resulting in insufficient sample sizes to make correlations. Each sample type displayed unique patterns of seasonal change across the land use types (Supp. Fig. R16) with seasonal fluctuations consistently observed only in endophytic Sphingomonadaceae. Sphingomonadaceae relative abundance in endophyte samples depended on its relative abundance in companion plants, but was shaped by a three-way interaction with season and land use (interaction Wald X^2^ = 15.0, p = 0.0006, Fig. 4C, Supp. Table R20). A negative correlation was observed in the fall for sites with strong anthropogenic influence, which changed to a positive correlation in the spring; we observed the opposite pattern for forest and railroad populations. Endophyte-soil (Supp. Fig. R17) did not display the three-way interaction for Sphingomonadaceae, but two-way interactions between land use and season explained significant variance (Fig. 4D, Supp. Table R20); no significant association was observed for soil-companion plant correlations. Overall, we observe that the strength and direction of microbial sharing differed across seasons and land use types at different levels of organization, from the macro-plant community to microbial diversity to Sphingomonadaceae relative abundance.

### Microbial abundance and diversity are associated with disease dynamics, but vary across land use types

Variance in disease PC1, which represents the number of symptoms, was better explained by bacterial than by fungal variation. Associations between bacterial load and disease PC1, depended on land use type and season (Fig. 5A, Supp. Table R21). As mentioned previously disease PC1 was higher in the spring (β = 0.27 ± 0.06 SE, p < 0.001, Fig. 5A). According to AIC, the best fit model included the interaction between disease PC1 and bacterial load (Supp. Table R22B), though we note that this interaction was not significant (two-way interaction Wald X^2^ = 5.1264, p = 0.07). For example, forest populations tended to have consistent bacterial loads across disease PC1, while anthropogenic and railroad populations were more variable. Additionally, disease PC1 was negatively correlated with bacterial Shannon diversity, with more symptoms observed in the plant associated with lower diversity (Fig. 5B). Companion plant community diversity still explained significant variance when accounting for bacterial effects on disease PC1 (Supp. Table R21). In contrast, disease PC2, which represents the symptom types, was better explained by variation in fungal microbiomes. Primarily, interactions between fungal load and-season explained significant variance for disease PC2 (interaction Wald X^2^ = 10.0, p = 0.002, Fig. 5C, Supp. Table R22), with a stronger and negative correlation in the fall.

**Fig. 5:**
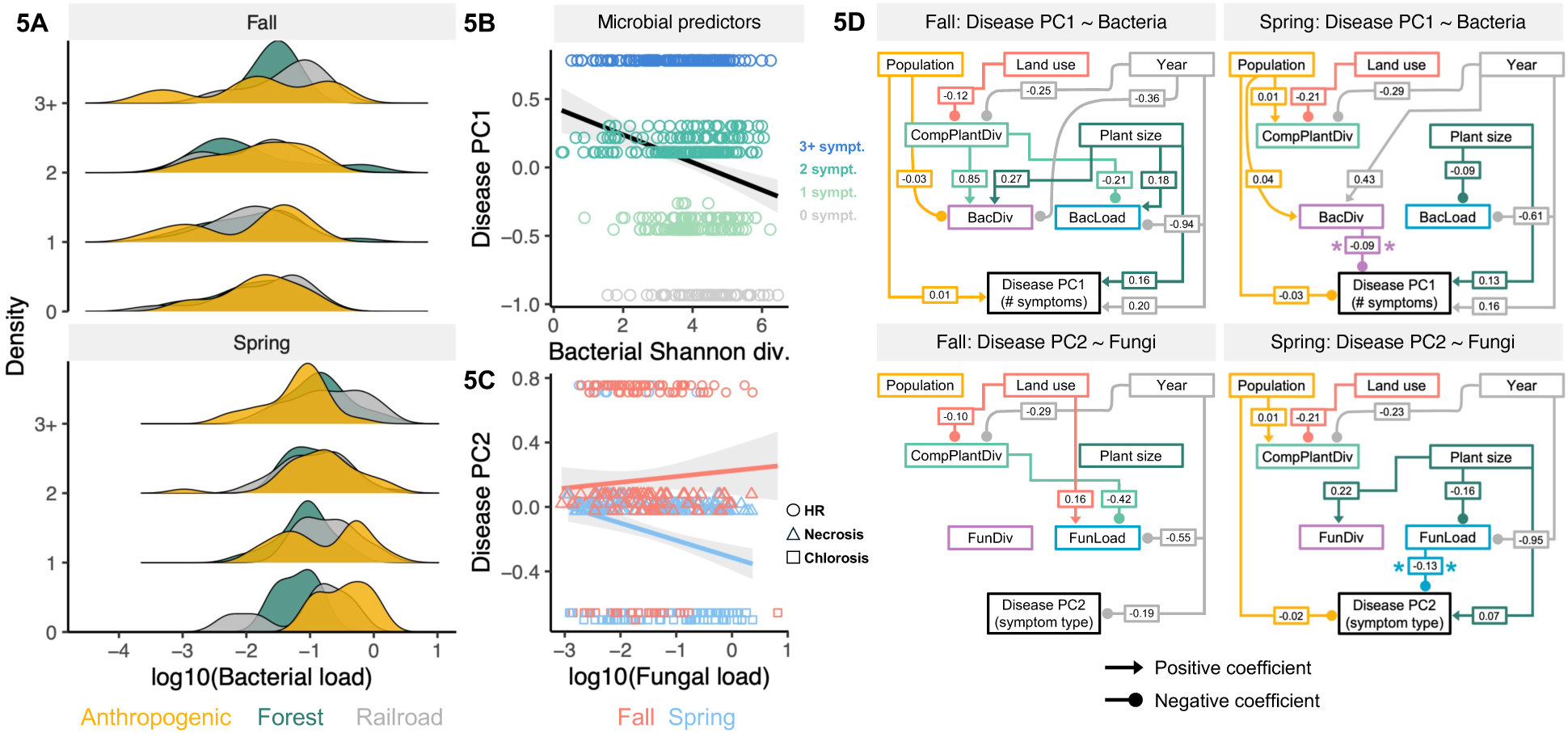
Associations between microbiome variation and disease symptoms vary by season, but differ between bacteria and fungi. A) Associations between bacterial load and disease PC1 as a function of land use type and season. Density plots reflect the number of individuals for a given disease PC1 (i.e., multiple-symptom class), colored by land use type. In B-C, we illustrate the negative association between disease PC1-bacterial diversity and disease PC2-fungal load. B) Association of Shannon diversity bacterial microbiomes with disease PC1 scores, with points colored by the number of symptoms observed. C) Association of fungal load with disease PC2, with points colored by season and shape reflecting symptom type associated with PC2 score. D) Path analysis results to explore hypotheses for how environment and microbiome contribute to variation in disease PC1 and PC2. Data were analyzed separately for each season to better visualize seasonal effects. Only in the spring did bacterial diversity directly impact disease PC1 and fungal load directly impact disease PC2. Only paths with significant coefficients (p < 0.05), were visualized, with the text boxes showing their coefficients. Direct paths between microbial traits in the spring are noted with asterisks.

We next performed path analysis to explore direct and indirect drivers that might explain significant variance in disease PC scores, given the complexity of the linear mixed model regressions as described in the previous paragraph. These paths represent hypotheses, and to better separate the interactions with season, we separated the analysis by season and between disease PC1 and PC2. While a complex set of interactions linked land use and companion plant diversity to microbial traits, only microbiomes were directly associated with disease PC1 (bacterial diversity) or PC2 (fungal load) in the spring (Fig. 5D).

Taken together, these analyses show the importance of considering multiple facets of microbial diversity (e.g., load and diversity from bacteria and fungi) to explain patterns in disease dynamics. Importantly, we find diversity at both the community and microbial level to be associated with reduced disease, highlighting how processes across ecological scales potentially impact disease dynamics.

## DISCUSSION

Here, we surveyed wild populations of *A. thaliana* across fall and spring seasons for disease symptoms and microbiome variation. While other studies typically focus on microbiomes or disease in the field, our study integrates both to show how land use impacts disease by modulating different aspects of the microbiome. Disease was primarily shaped by interactions between season and land use. In the spring, more individuals display multiple disease symptoms, with higher microbial load and changes to microbial community composition (Figs. 2-3). Increased companion plant community diversity was associated with reduced occurrence of multiple disease symptoms, but this relationship differed between land use types (Fig. 4). Intriguingly, plant community diversity was positively associated with increased *A. thaliana* microbial diversity (Fig.4), and this higher microbial diversity was also associated with reduced disease in *A. thaliana* (Fig. 5). Our results highlight how land use type can impact multiple modes of ecological interactions that in turn can change disease risk in wild populations.

We observed that higher biodiversity at two ecological scales–the plant community and within the *A. thaliana* microbiome–resulted in less disease. This result is suggestive of the dilution effect, where higher host diversity dilutes the disease risk in a multi-host pathogen system [1,8,58]. Biodiversity loss associated with urban and agricultural environments often results in communities enriched for species with life history traits associated with fast pace-of-life and reduced investment in immune defenses, increasing the risk of disease spillover [3,58]. While we did not measure these traits in the companion plant communities, land use types differed in the correlation between disease symptoms and plant diversity (Fig. 2D) as well as plant community composition (Supp. Fig. R3), suggesting that traits underlying the diversity-disease relationship vary across land use types. The impact of this variation in disease-relevant plant traits can also be further modulated by the abiotic factors, such as soil chemical properties [41]. Higher diversity microbiomes often provide greater protection against pathogens in plants [15,17,59], as we observed here as well, but this protection also partially depends on microbial load (Fig. 5). Protection by diverse microbiomes occurs when commensals saturate the host niche space and consume resources that effectively outcompete invading pathogens [60,61]. Beyond just competition over resources, microbiomes can also directly inhibit other microbes through the production of antimicrobial agents [20] or indirectly by priming the plant immune system [21,22]. Identifying potential synergies between the mechanisms underlying the dilution effect at the plant community scale and within the microbiome is an important research priority.

Our results are consistent with several scenarios, but we believe that the following is particularly attractive. First, plant community diversity protects against disease through plant traits that dilute the community-level disease burden. In turn, increases in plant community diversity also promote diversity within the *A. thaliana* microbiome, resulting in increased protection and reduced disease. Our path analysis (Fig. 5D) to some extent supports a scenario whereby land use reshapes companion plant diversity, which in turn reshapes microbial diversity and load. Although these effects are observed in both the fall and spring, bacterial diversity and fungal load were only directly associated with disease dynamics in the spring. These results suggest that there are seasonal differences in the relationship between companion plant communities and microbiome dynamics that may also impact disease dynamics, but this is largely hypothetical. While it is difficult to know why effects of plant diversity were not observed in the fall, the pattern is consistent with transitory dynamics dominating as the microbiome community becomes established [62]. One potential issue is that we hypothesized a specific directionality in our path analysis, but that there could be unaccounted feedbacks within these scenarios. Additional support is required to untangle these potential outcomes, preferably from targeted manipulation of microbiomes, pathogens and companion plants. We note that the dilution effect often depends on the spatial scale over which these interactions occur [63,64]; it remains an open question how to apply the framework of the dilution effect to host-associated microbiomes [1,63].

The concomitant increase in diversity between plant community and microbial diversity with increased microbial diversity in *A. thaliana* exposes a tension in the diversity-disease relationship. This diversity-begets-diversity relationship has been observed in a range of host systems, including *A. thaliana* [26,58,65], but primarily for pathogens rather than commensal microbes. The positive relationship between plant community and microbial diversity appeared to be beneficial in our setting, but it also potentially amplifies disease risk by exposure to novel microbes. Microbial sharing between host species is expected to accelerate microbial evolution particularly under environmental change [14,66], but whether cross-species transmission differentially impacts pathogens compared to commensal microbes is an open question. Intriguingly, the correlation of Sphingomonadaceae between *A. thaliana* endophyte samples and the other sample types suggests that land use alters microbial sharing (Fig. 4C-D). Sphingomonadaceae abundance is often protective against bacterial pathogens [21,67], but as we did not assess disease in companion plants, we do not know if this occurred in the surveyed plants. A key research priority is to link how these different eco-evolutionary forces shape the microbial evolution in wild host-microbe systems.

While it is tempting to attribute the negative relationship between microbial diversity and disease to protection provided by the microbiome or the dilution effect operating at the microbial scale, we do not know microbiome diversity prior to infection and thus cannot attribute causality. Furthermore, we failed to detect strong effects of disease on structuring the beta-diversity of the endophytic microbiome (Supp. Fig. D1), suggesting that microbial taxa did not consistently differ between healthy and sick individuals. One issue is that pathogens often share the same 16S rDNA sequence as non-pathogenic microbes in plant disease systems, such as in the *P. syringae* complex [44], and so we have limited insight from our amplicon sequencing to identify pathogens. We attempted to use the *HSD* amplicon for *Pseudomonas* genera, but we did not find any associations with the presence of HR-like lesions or the total number of disease symptoms in diseased plants (Supp. Fig. D2), suggesting this amplicon lacks sufficient resolution to identify pathogenic strains. While there may be technical limitations, Shannon diversity was the primary difference for the microbiome between healthy and diseased plants, leaving much of the variation in disease-microbiome interactions unexplained.

It is remarkable that so many *A. thaliana* individuals were observed to have multiple disease symptoms, and understanding this resilience of *A. thaliana* to disease pressure remains a key question. While multi-pathogen systems are common in the wild ecosystems, individuals seem to rarely experience strong negative effects of coinfection [23,24], which is aligned with our observations here. Additionally, infections did not appear to cost *A. thaliana* in growth or reproduction, as larger plants tended to have more disease symptoms and were just as likely to be bolting as healthy plants (Supp. Fig. D3). Note, however, that we did not measure phenotypes more directly associated with fitness. We also do not know when plants had been infected or if they had cleared prior infections. It is possible that disease symptoms may not necessarily be independent, as the same pathogen can cause multiple symptom types [68]. We cannot confirm the association between the number of symptoms, type of symptoms, and the number of pathogens due to the lack of resolution in amplicon sequencing.

This lack of resolution applies to both bacterial and fungal microbiomes. As previously mentioned, neither 16S rDNA nor *HSD* amplicons unequivocally identified potential pathogens. Other amplicons, like *gyrase B*, can identify pathogenic bacteria within the plant microbiome [26] and their use would be an important next step in the type of work we describe here. Particularly for fungal microbiomes, the ITS1-2 region provides limited insight due to high intra- and intergenomic variation in fungal genomes, although it remains the best option for fungal metabarcoding [69]. Annotated databases, like FunGuild [52], can help link ITS variation to function, but their utility depends on the training data. We classified the trophic mode for the fungal genera observed here, and most were simultaneously classified as pathotroph, saprotroph, and symbiotroph (Supp. Fig. D4). The assignments of detected fungi also did not vary across disease states, providing limited insight into potential causative agents of the variation in symptoms observed. In addition, the symptoms observed here may not be due solely to disease, but may also reflect exposure to environmental stress. Diagnosing the causative agents of symptoms combined with transcriptomic profiling would provide clarity into the relationship between land use and disease in *A. thaliana*.

Finally, it is important to consider the role of plant genetics and demography in shaping interactions between pathogens and the microbiome. Host genetics impacts microbiome composition in *A. thaliana* [28–30,33,70] as well disease resistance [29,71–73]. Land use in turn impacts host genetics, as different urban environments filter for different *A. thaliana* genotypes [74]. *Arabidopsis thaliana* has been shown to locally adapt to the companion plant community, particularly for alleles in immunity genes [40]. Railroads are implicated in increasing gene flow in a closely related species, *A. arenosa* [75]. Whether this applies to the *A. thaliana* populations here remains unknown, especially whether the introduced alleles by railroad-assisted gene flow would be associated with host control over the microbiome or disease resistance in the populations sampled. Both studies of urban adaptation [74] and introgression facilitated by railroads [75] identified selection on alleles associated with life-history traits that shorten the life cycle. Changes to the demography of *A. thaliana* across different land use types may lead to multiple cohorts of germination during each growing season. If these germination cohorts differ from the temporal dynamics of pathogen exposure, then this may explain seasonal differences in both the individual symptoms (Supp. Fig. R4) and composite disease scores (Fig. 2C) observed across land use types. Repeated sampling at smaller time intervals would help elucidate how temporal variation in pathogen exposure intersects with *A. thaliana* genetic variation and demography.

In conclusion, our results highlight the multitude of ways that land use affects ecological interactions, from the macro-community level down to within-host microbial variation to plant disease dynamics. For the many ecological communities impacted by rapid anthropogenic change, our work here highlights the importance of understanding the interaction between seasonality and ecological context that will shape disease dynamics in natural populations. Harnessing these interactions will be critical in developing the microbiome as a potential solution to provide protection against pathogens in wild and agricultural settings.

## ACKNOWLEDGEMENTS

We thank L. Burkhardt, A. Contreras, S. Dambrowski, L. van Ess, K. Fritschi, A. Habring, C. Lino, M. Lucke, D. Lundberg, A. Marotz, E. Mehmetoğlu, A. Nowakowski, G. Ofir, K. Romanova, R. Schwab, S. Stegmeyer, P. Sterr, L. Teasdale, S. Wang, F. Weber for assistance in the field and lab. We acknowledge the Zegar Family Foundation for their generous support of the NYU Genomics Core. LPH was supported by the Charles H. Revson Foundation. This work was funded by grant SFI-PD-Grant-01308072 of the Simons Foundation to JB and ERC-SyG Project 951444 (PATHOCOM) awarded to FR, DW, and JB.

## DATA AVAILABILITY

Data are available on [Zenodo](https://zenodo.org/records/15769719?token=eyJhbGciOiJIUzUxMiJ9.eyJpZCI6IjUzMzE0M2MyLTA4NjctNDZiNS1hYjE5LTI1MWYxNjYxNDZlNSIsImRhdGEiOnt9LCJyYW5kb20iOiI1ZTgwMWRjNDQyNmY0ZmRlNjkzMWFjYTU4ZWZjZjQ2ZiJ9.uxV1f8rtnKzTjqqI_Y-eCY041FlN06IlL79hlk9n01XF-bd1YAN6_t4S0D4lo81LZAJVZrX80pCUFKKJEzLIpQ). Code is available on [Zenodo here](https://zenodo.org/records/15769773?token=eyJhbGciOiJIUzUxMiJ9.eyJpZCI6IjI4NWY0NDQ0LWU4MDUtNDUyZC1iY2Q3LTcwMzBjNDk3OGVkMiIsImRhdGEiOnt9LCJyYW5kb20iOiJlMzJjZDY2YjdmZTk3ZmQzMWViNDE5NDIzMWQ5NWE4MyJ9.imYB4i4REw12RJ9vUkt0dmZi5EukH-NHiiDklqpYcneXcbL10dJ_96xMn0xZpoQoLcKFivRavCmpu32f3fOmfQ). Sequencing data are uploaded to NCBI SRA, as Bioprojects PRJNA1279102 for 16S rDNA, PRJNA1282642 for ITS1-2, PRJNA1282654 for HSD, and PRJNA1282661 for mmdA.

