## supplemental information for "Seasonality, land use, and host diversity shape microbiome-pathogen interactions in wild populations of *Arabidopsis thaliana*"

### ***SUPPLEMENTAL METHODS***

#### *Sampling A. thaliana*

Sites with *A. thaliana* populations were identified across a variety of land use types in Michigan, USA. Sampling occurred once per fall and spring from Fall 2021 through Spring 2023. At each site, two transects were established that captured the range of environmental variation. The GPS coordinates of each transect were recorded and positions marked in the sites to enable us to return to approximately the same transects over the next two years. Quadrats for characterizing the companion plant communities (see below) were placed in the middle of the transect, but were sometimes moved slightly to ensure at least one *A. thaliana* individual was located within the quadrat. As mentioned in the main text, the transects were pooled together for all analyses.

While we attempted to collect at all sites during the two years of sampling, there were several exceptions. Site 17 was only visited in Fall 2021, but we did not return to this site in subsequent seasons. Site 19 was added to make up for this lost site, and thus sampling began in the Spring 2022. In Fall 2022, no *A. thaliana* plants were identified at Site 4 and thus we did not sample. Sites 6, 7, and 16 were covered in ~15 cm of snow in Fall 2022 and were not sampled.

At each site, we first estimated the *A. thaliana* population size. Then, we identified 16-24 *A. thaliana* individuals across the two transects, and collected only their rosettes. Rosette diameter was measured, number of leaves counted, bolting status, and then scored for disease symptoms (chlorosis, necrosis, hypersensitive response (HR)-like lesions, *Albugo* presence, *Hyaloperonospora* presence, general fungal damage), and herbivory. HR-like lesions were classified as presence/absence, but the other symptoms were initially scored on a scale of 0-4 (0 = absence of symptoms, 4 = severe symptoms). For all analyses presented here, disease symptoms were collapsed to presence/absence.

After plants were phenotyped, approximately half were selected for microbiome profiling. These plants were processed in the field to enrich for the endophytic portion of the microbiome as follows. Care was taken to minimize contamination by cleaning all tools with bleach and 70% ethanol; all reagents were sterilized before bringing into the field. First, any soil, roots, or residual organic matter were gently removed with forceps. Plants were placed into a tube with sterile water and inverted 10 times. Rosettes were then transferred to another tube with 1X TE + 0.01% Triton-X and inverted 10 times. This was repeated in a fresh TE+Triton-X tube. Then the rosette was transferred to a tube with 2% bleach and inverted 10 times, followed by two washes with sterile water. The rosette was blotted dry, placed into a new tube, stored on dry ice in the field, and at -80ºC upon returning to the lab.

Companion plant and soil samples were also collected for microbiome profiling, please see below for these procedures.

#### *Soil collection*

Soil samples were collected for microbiome profiling at each site and sampling time point. Four soil samples per site were collected into 2 ml tubes from locations adjacent to and within the quadrat. DNA was extracted and libraries prepared from each individually, but then the data was pooled per site.

For characterizing the physicochemical properties, soil samples were collected at each site during the Fall 2021 and Spring 2022 of the first sampling season. Approximately 1 kg of soil was collected. Soil was stored at 4ºC in the lab and then was analyzed at A&L GreatLakes

Laboratories for 20 chemical properties using standard methods (https://algreatlakes.com/pages/primary-method-references-soil). Soil properties were averaged together between fall and spring. To reduce the number of covariates in subsequent analyses, we performed principal component analysis (PCA) using prcomp and factoextra in R. Soil properties were scaled prior to PCA, and the first two principal components were used.

*Characterizing the companion plant community*
As described in the main text, two 50x50-cm quadrats were placed in the approximate center of each transect, in areas with at least one *A. thaliana* positioned within the quadrat. Each quadrat was subdivided into 25 smaller equal sized 10x10-cm squares. We first identified morphotypes and as proxies for plant species and counted each morphotype per each of the 25 small squares. Per each small square, 10-20 individuals for a given morphotype were counted as "10" and greater than 20 were counted as "20". A voucher for each morphotype was collected per transect for molecular identification (more below). If there was any uncertainty about the similarity between two morphotypes, these were counted as distinct morphotypes, each saved as a voucher to later confirm the identity, and then summed together when appropriate. For all analyses, we pooled together the transects per site.

Two samples of the companion plant communities were collected for microbiome profiling at each site and sampling time point form each quadrat. Portions of leaves from each morphotype were collected and placed into a sterile 2 ml tube. DNA was extracted and libraries prepared from each individually, but then the data was pooled per site.

To confirm plant identity, we used a metabarcoding approach with the chloroplast marker *matK* as previously described [1]. In brief, DNA was extracted as described below, and then a cocktail of PCR primers were used to amplify *matK* (Supp. Table S1). *matK* sequences were initially clustered into the OTU level at 98% similarity using USEARCH [2], and taxonomy was assigned using the NCBI BLAST Nucleotide Collection. We identified 122 OTUs across 63 genera, and we used these genus-level assignments in our analyses. Out of the ~33,000 companion plants counted in our surveys, we were could classify 98.2%. Unclassified plants were not considered for alpha- and beta-diversity analyses.

*Temperature and precipitation data*
Temperature was recorded over the two years of sampling at each site using an iButton temperature logger (https://www.ibuttonlink.com/). Two iButtons were placed per site near transects, but often under cover of bushes or other shading to hide them within the sites. Because these iButtons were not precisely paired with each transect, we averaged together the temperature readings per site. At each sampling (e.g., Fall 2021, Spring 2022, etc.), we downloaded the temperature recordings and returned iButtons to their location within the site. The mean temperature on the day of sampling was used in subsequent analyses.

While the majority of temperature data was collected using iButtons, we note that some information was lost due to missing or malfunctioning iButtons. In this case, we supplemented the data with publicly available weather data (https://www.wunderground.com/). We identified the closest weather station to the sites and collected the available data for the three weeks preceding each sampling (Supp. Table S2 for list of weather stations). We note that some sites were covered by the same weather station. Temperature measurements were correlated between iButtons and Weather Underground ($F_{1,43} = 68.26$, $p < 0.0001$, $R^2 = 0.61$, Supp. Fig. S1).

Precipitation data was collected from the same weather stations using Weather Underground. Precipitation amounts were summed together from the three weeks preceding the collection for each season. This cumulative precipitation measure was used in subsequent analyses.

*DNA extraction and library preparation*
DNA extraction was performed as described [3]. In brief, either ~0.5 ml of garnet rocks or a mix of silica beads (ranging from 0.5 mm to 3.0 mm) were added to tubes and samples were initially ground under liquid nitrogen. Then 800 µl of lysis buffer (10 mM Tris pH 8.0, 10 mM EDTA, 100 mM NaCl, 1.5% SDS, 40 mM dithiothreitol, 20 mg/ml proteinase K) was added and homogenized for 1 min on a Spex Geno/Grinder at max speed. Samples were incubated at 65ºC for 1 hour. Plant debris was pelleted and then 400 µl of lysate was added to 150 µl potassium acetate to precipitate the SDS. Samples were centrifuged twice and the debris-free supernatant was mixed with SPRI beads at a ratio of 0.58. After mixing and incubation, beads were washed twice with 80% ethanol and eluted in sterile water. DNA was quantified using PicoGreen assay and normalized to ~10 ng/µl. If DNA concentration was below 10 ng/µl, DNA was not diluted. All plates contained at least 2 process blanks to use later to assess for potential contamination in microbial analyses.

To profile the bacterial and fungal microbiomes, libraries were prepared using the host-associated microbe PCR (hamPCR) method as previously described [4]. hamPCR follows a standard two-step, dual indexed PCR procedure that amplifies the genes of interest in the first PCR and adds Illumina adapters in the second PCR. In the first PCR, primers for the microbial genes(16S rRNA V5V6V67 or ITS1-2) and host genes (*GIGANTEA*) were mixed equally at 10 µM concentrations, and then 0.4 µl of the primer mix was used per 10 µl reaction. The rest of the PCR reaction contained 5 µl HotStart OneTaq (NEB M0481), 2 µl DNA, and sterile water to 10 µl. Cycling conditions in the first PCR for both 16S and ITS were: 98ºC for the initial denaturation of 3 min, 10 cycles of 98ºC for 30 s, 59ºC for 30 s, 68ºC for 30s, and 68ºC for 5 min in the final extension. After the first PCR, ExoSap purification was performed to remove un-annealed primers. The second PCR added Illumina adapters with the following PCR conditions:  98ºC for the initial denaturation of 3 min, 30 cycles of 98ºC for 30 s, 55ºC for 30 s, 68ºC for 30s, and 68ºC for 5 min in the final extension. Libraries were pooled per PCR plate using equal volumes and cleaned using SPRI beads. For final cleaning,  ~600 bp were excised and cleaned up for sequencing using the Zymoclean Gel DNA Recovery kit (Zymo D4008). Quality and quantity of pools were assessed using TapeStation D1000 and then pooled equimolarly for sequencing on Illumina platforms.

There were differences in the library preps for *A. thaliana* endophyte and the other sample types. First, *GIGANTEA* primers were only added to the *A. thaliana* endophyte samples. Second, 0.2 µl of 50 mg/ml UltraPure bovine serum albumin (BSA, ThermoFisher AM2616) per 10 µl PCR reaction were added to only companion plant and soil PCRs, as these sample types often have many secondary compounds that inhibit PCR [5]. Third, to minimize biases that occur during the logarithmic amplification in the first PCR for hamPCR, *A. thaliana* endophyte samples were only amplified for 10 cycles in the first PCR. Soil and companion plant samples were amplified for 30 cycles in the first PCR and 10 cycles in second PCR. hamPCR has previously been shown to be robust to the cycle number variation [4], and we further confirmed that 10 or 30 cycles did not distort our assessment of soil and companion plant microbiome composition (Supp. Fig. S2).

Libraries were sequenced on the Illumina NovaSeq 6000 platform at the NYU Genomics Core. For both 16S and ITS libraries, most endophyte samples were sequenced on two runs to increase the microbial read depth, and reads were pooled together from the sequencing runs. Most soil and companion plant samples for 16S rDNA libraries were prepared using both 10 cycles and 30 cycles in the first PCR. As there was no effect on community composition, reads were combined.

To profile strain level diversity within *Sphingomonas* and *Pseudomonas* genera, we did not use hamPCR, but the standard two-step, dual indexed amplicon library procedure. Please see the section “Designing *mmdA* and *HSD* amplicons” for details on primer design. The first PCR 98ºC for the initial denaturation of 3 min, 25 cycles of 98ºC for 30 s, 60ºC for 30 s, 72ºC for 1 min, and 72ºC for 10 min in the final extension. The second PCR added Illumina adapters with the following PCR conditions: 98ºC for the initial denaturation of 3 min, 15 cycles of 98ºC for 30 s, 55ºC for 30 s, 68ºC for 30s, and 68ºC for 5 min in the final extension. Libraries were pooled per PCR plate using equal volumes and cleaned using SPRI beads. For final cleaning, ~600 bp were excised and cleaned up for sequencing using the Zymoclean Gel DNA Recovery kit (Zymo D4008). Quality and quantity of pools were assessed using TapeStation D1000 and then pooled equimolarly for sequencing on the Illumina MiSeq platform.

*Design of mmdA and HSD primers*
To create primers to distinguish different strains within a clade we used the following approach. First, for a set of genomes from a given clade we predicted all proteins with prodigal [6]. We then used diamond [7] with default settings to identify gene families. We considered a gene family a core gene if it was present in at least 95% of the genomes, and we only considered core genes which were present at a single copy in at least 98% of the genomes in which they occurred for further analyses.

We created codon alignments for each core gene by aligning the protein sequences using mafft [8] and a custom python script to create the codon alignments. We ran emboss cons [9] to create a consensus sequence for each codon alignment, using an identity cutoff of 0.95. That is, all positions in the alignment for which one base was not present at least 95% were considered Ns in the final alignment. Then for each single-copy core gene we ran primer3 [10] with parameters specified in the custom python script "primer3_file_writer.py."

Finally, to evaluate candidate primers we used a custom python script which took a given primer pair and extracted all the amplicons that would be amplified by this primer pair across the genomes. Then we sorted the primer pairs by the number of unique sequences to identify candidate primer pairs for further testing by PCR. For these primers, we converted all N bases to the appropriate degenerate bases using a custom python script. This resulted in the following primers pairs:

mmdA-forward: CGGTTCGTSCGCTTCT
mmdA-reverse: TTGTCATGCTTCTTCCACGG
HSD-forward: TYCCGGCYGAYCGYCTGATYGCCA
HSD-reverse: TGVGTCARCAGRATCATBGGCACCW

We validated the ability of different primer pairs to produce single bands from both pure bacterial genomic DNA and wild plant microbiome samples, and not to produce any bands from pure plant genomic DNA.

*Bioinformatic analyses*
The reads were basecalled using Picard IlluminaBasecallsToFastq version 2.23.8[11], with APPLY_EAMSS_FILTER set to false. Reads were demultiplexed using barcode_splitter [12]. Illumina adapters and polyG sequences were trimmed using fastp v0.20.1 [13]; reads with fewer than 250 bp were discarded after this filtering. Because each endophyte library contained both microbial and host genes, before calling ASVs, we split the libraries into microbial or host *GIGANTEA* reads using bbtools v38.91 [14]. Soil and companion plant libraries contained spurious and low amounts (<0.1%) of *GIGANTEA* reads, we also used bbtools to separate plant libraries to ensure that libraries were computationally processed the same way as the endophyte samples.

QIIME2 v2023.2 [15] was used to call ASVs. Reads were quality filtered and trimmed to 200 bp for 16S rDNA and its paired *GIGANTEA* reads, and 250 bp for ITS and its paired *GIGANTEA* reads. DADA2 [16] was used to call ASVs. Taxonomy was assigned using classify-sklearn Bayes Classifier [17]. 16S rDNA sequences were classified using the Greengenes database [18] and ITS sequences were classified using the Unite database [19]. After taxonomic assignment, all reads that were not classified at the phylum level were discarded. We used decontam [20] to filter out ASVs associated with our negative controls at threshold of 0.2. After filtering potential contaminants, if a sample was sequenced more than once, all reads were summed together.

For microbial load, only microbial and *GIGANTEA* reads that were successfully called as ASVs were included in the calculation. This approach excluded anything that was not successfully sequenced (e.g., adapter read through or was determined to be a PCR chimera). Microbial load was the total number of microbial reads divided by the *GIGANTEA* reads. For microbial reads, only ASVs that were assigned at least the phyla level were included.

For beta- and alpha-diversity analyses, we filtered out any ASVs not seen in at least 2 samples and ASVs below <0.1%. Beta-diversity in the microbiome was based on a modified Aitchison distance matrix [21]. While this distance metric is largely insensitive to read depth, we included read depth as a covariate, as rarefaction was not performed before ordination. For alpha-diversity in the microbiome, we rarefied to 1,000 reads for each of the four amplicon types in endophyte samples (Supp. Fig. M3 for rarefaction curves) and then calculated alpha-diversity measures. We rarefied to 16,963 reads for companion plant and soil microbiomes. Our justification for the two different depths is that reduced diversity was observed in endophyte samples, the saturation point in the rarefaction curve was lower, and this enabled us to retain most endophyte samples. We had much higher read depth for the soil and companion plant microbiomes, and so we rarefied down to the lowest read depth per sample. As endophyte samples were also related to individuals, while the companion plant/soil microbiomes were representative per site and sampling season, this further justifies using different read depths between endophyte and the other sample types.

*Statistical analyses*
We outlined the general statistical approach for all analyses in the main text. As a brief recap, random effects per population was nested within year (denoted as 1|Year:Population). We describe how interactions between terms were handled for each test, but generally non-significant interactions were dropped. Next, we describe in detail each statistical test performed, and where applicable, we reference the figure that displays the data and table that summarizes the result. Some variables were transformed to ensure normality of residuals, and the different transformations are denoted in each model.

Testing for effects of land use on *A. thaliana* demographics (population size, plant size):
The model to test for differences in *A. thaliana* population size and land use type (Fig. 1B, Supp. Table R1), is below. The interaction between land use and season was not significant.

log10(population size) ~ Land use + Season + (1|Year:Population)

The model to test for differences in plant size and land use type (Fig. 1C, Supp. Table R2) is below. The interaction between land use and season was not significant. Sqrt = square root transformation.

sqrt(Max diameter) ~ Land use + Season + (1|Year:Population)

Testing for effects of land use on weather data:
The model to test for differences in temperature profiles between land use types (Supp. Fig. R1, Supp. Table R3) is below.

Temperature ~ Land use + latitude + (1|Population)

The model to test for differences in precipitation between land use types (Supp. Fig. R1, Supp. Table R3) is below. The interaction between land use and season was not significant.

Precipitation ~ Land use + Season + (1|Year:Population)

Testing for effects of land use on companion plant community:
The model to test for factors explaining significant variation in percent coverage by the companion plant community (Supp. Fig. R3, Supp. Table R4) is below. The interaction between land use and season was not significant. Sqrt = square root transformation.

sqrt(Coverage) ~ Land use + Season + Soil PC1 + Soil PC2 + (1|Year:Population)

For analysis of beta-diversity in the companion plant community (Supp. Fig. R3, Supp. Table R5), we used Bray-Curtis distance and PERMANOVA blocked by Population. The significance was assessed by each term in the sequential order of the model below:

Bray Curtis ~ Year + Season + Land use + Soil PC1 + Soil PC2 + Temperature + Precipitation

For alpha-diversity analyses in the companion plant community (Supp. Fig. R4, Supp. Table R6), we first calculated three measures of alpha-diversity: Observed richness, Shannon diversity, and Evenness. For each measure, the statistical model is below. The interaction between land use and season was not significant for any of the alpha-diversity measures.

Alpha-diversity measure ~ Land use + Season + (1|Year:Population)

Testing for effects of land use on disease symptoms:
To test for factors that explained significant variation in the disease scores (Fig. 2, Supp. Table. R7), we focused on the disease PC1 as this separated multiple-symptoms. Because we were

comparing two models, both were fitted using the maximum likelihood method. We first fitted the model to test for interactions between land use type and season:

Disease PC1 ~ Land use * Season + Max_diam + (1|Year:Population)

We then tested if including companion plant diversity explained significant variation by adding this term as follows:

Disease PC1 ~ Land use * Season * Companion plant Shannon + max_diam + (1|Year:Population)

The interaction terms in both models were significant. We then used the likelihood ratio test to identify which model explained significantly more variation. The significance of the three-way interaction terms was assessed using type III Wald $X^2$ tests.

Testing for effects of land use on the microbiome

To test for drivers of microbial load (Fig. 3A-B, Supp. Table R8), we separately fitted models for bacterial and fungal load. The interaction between land use and season was not significant.

log10(microbial load) ~ Season + Land use + max_diam + (1|Year:Population)

To test if bacterial and fungal loads were correlated (Supp. Fig. R7, Supp. Table R9), we fitted the model below. The significance of the two-way interaction was assessed using type II Wald $X^2$ tests.

log10(bacterial load) ~ log10(fungal load) * Season + max_diam + (1|Year:Population)

To test for the contribution of different factors explaining variance in composition between sample types (Supp. Fig. R10, Supp. Table R10), we used a modified Aitchison distance matrix and PERMANOVA blocked by Population. Models for bacterial and fungal microbiomes were tested separately, but with the same terms. The significance was assessed by each term in the sequential order of the model below:

Aitchison distance ~ Sample type + Season + Year + Land use + log10(reads)

We confirmed that similar qualitative results were observed using two alternative distance metrics, Bray-Curtis and weighted UniFrac (Supp. Fig R11, Supp. Table R11). For both Bray-Curtis and weighted UniFrac, samples were rarified down to 1000 reads as these distance metrics are more sensitive to read depth than the modified Aitchison distance [21]. The same models were run as above, but without the log10(reads) term.

We compared differences in alpha-diversity metrics for the different sample types (Supp. Fig. R12, Supp. Table R12) using the Kruskal-Wallis test statistic because the variance was very different between each sample type. ASV richness, Faith's phylogenetic distance, evenness, and Shannon diversity were all compared between the three sample types.

We then focused on the comparisons of forces structuring beta-diversity within each sample type. We included the quantitative variables (e.g., soil chemistry PC1, temperature, etc) in these analyses to more specifically identify structuring forces between the different sample types. The models were the same for both bacterial and fungal microbiomes across the sample types. We tested for the different factors explaining variance in composition between sample types (Fig. 3, Supp. Table R13 for bacteria, Supp. Table R14 for fungi), using a modified Aitchison distance matrix and PERMANOVA blocked by Population. The significance was assessed by each term in the sequential order of the model below:

Aitchison distance ~ Season + Year + Land use + Soil chemistry PC1 + Companion plant beta-diversity PCoA1 + Mean temperature + Precipitation + log10(reads)

We examined how Shannon diversity in different amplicons (*16S, ITS, mmdA, HSD*), which reflect different phylogenetic scales of diversity, changed over time and across land use types (Fig. 3G, Supp. Table R15). The model was used below. The significance of the three-way interaction was assessed using type III Wald $X^2$ tests.

Shannon diversity ~ Amplicon * Season * Land use + max_diam + (1|Year:Population)

To test for factors that drive Shannon diversity differences within companion plant and soil microbiomes (Supp. Fig. R13, Supp. Table R16), we fitted the statistical model below. The interaction between land use and season was not significant.

Shannon diversity ~ Land use + Season + (1|Year:Population)

To test for correlations between Shannon diversity and microbial load Supp. Fig. R14, Supp. Table R17), we fitted the model below. For bacterial load, the significance of the two-way interaction was assessed using type II Wald $X^2$ tests. There was no significant interaction for fungal load, and thus the interaction term was dropped from the model.

Microbial Shannon diversity ~ log10(microbial load) * Land use + Season + max_diam + (1|Year:Population)

To test for correlations between companion plant community diversity and endophyte microbial diversity (Fig. 4A-B, Supp. Table R18), we fitted the model below for both bacterial and fungal microbiomes. The significance of the two-way interaction was assessed using type II Wald $X^2$ tests. Three-way interactions between companion plant community diversity, land use, and season were not significant.

Microbial Shannon diversity ~ Companion plant community Shannon diversity * Land use + season + max_diam + (1|Year:Population)

To test for correlations between microbial diversity in the endophytes with microbial diversity in the soil and companion plant (Supp. Fig. R15, Supp. Table R19), we fitted the model below for bacterial and fungal microbiomes. We started with three-way interactions for all models, but dropped the interactions if not significant. We used the appropriate Wald $X^2$ test for the number of interactions. We did not test correlations between bacterial and fungal microbiomes. Each model is described below:

16S Endophyte Shannon ~ 16S companion plant Shannon * Land use + Season + max_diam + (1|Year:Population)

ITS Endophyte Shannon ~ ITS companion plant Shannon * Land use * Season + max_diam + (1|Year:Population)

16S Endophyte Shannon ~ 16S soil Shannon + Land use + Season + max_diam + (1|Year:Population)

16S Endophyte Shannon ~ 16S companion plant Shannon * Land use + Season + max_diam + (1|Year:Population)

ITS Endophyte Shannon ~ ITS soil Shannon * Land use * Season + max_diam + (1|Year:Population)

To test for microbial sharing, we asked whether Sphingomonadaceae relative abundance in endophytes was predicted by abundance in the environmental reservoirs of the companion plant and soil microbiome (Fig. 4C-D, Supp. Fig. R17, Supp. Table R20). For each sample type comparison (e.g., endophyte-companion plants), we first fitted the following model for each land use type and season to estimate the correlation coefficient for the endophyte-reservoir association. Sqrt = square root transformation.

sqrt(rel. abund. endophyte) ~ sqrt(rel. abund. reservoir) + (1|Year:Population)

Then, to formally test for factors that shape Sphingomonadaceae sharing, we fitted the model below for endophyte-companion plant associations. The significance of the three-way interaction was assessed using type III Wald $X^2$ tests.

sqrt(rel.abund. endophyte) ~ sqrt(rel. abund. companion plant) * Land use * Season + (1|Year:Population)

However, three-way interactions were only observed in endophyte-companion plant comparisons. The endophyte-soil association was as follows, and significance was assessed using type II Wald $X^2$ tests.

sqrt(rel. abund. endophyte) ~ sqrt(rel. abund. soil) * Land use + Season + (1|Year:Population)

The soil-companion plant association was assessed as follows, as no interactions were significant:

sqrt(rel. abund. companion plant) ~ sqrt(rel. abund. soil) + Land use + Season + (1|Year:Population)

<u>Testing for associations between the microbiome and disease symptoms</u>
We tested for the association of different microbial parameters in the *A. thaliana* endophytes and disease symptoms using a model selection approach. We included both microbial load and Shannon diversity to identify the factors that explained the most variance. We fit with 16S and ITS data separately as load was positively correlated between bacterial and fungal microbiomes (Supp. Fig. R7). We fitted models with microbial load and diversity separately as they were correlated with each other (Supp. Fig. R14). For each model, we assessed the significance of interaction terms using Wald $X^2$ tests starting with three-way interactions, dropping non-significant interactions, and then the likelihood ratio test was used to find the best fitted models. We made the comparisons separately for disease PC1 and PC2.

For disease PC1 (Fig. 5, Supp. Table R21), the models were as follows:

M1: disease PC1 ~ log10(bacterial load) * Land use + season + max_diam + (1|Year:Population)

M2: disease PC1 ~ Bacterial Shannon diversity + Land use + season + max_diam + (1|Year:Population)

M3: disease PC1 ~ log10(bacterial load) * Land use + Companion plant community Shannon diversity + season + max_diam + (1|Year:Population)

M4: disease PC1 ~ log10(fungal load) + Land use + season + max_diam + (1|Year:Population)

M5: disease PC1 ~ Fungal Shannon diversity * Land use * season + max_diam + (1|Year:Population)

M6: disease PC1 ~ log10(bacterial load) * Land use + Companion plant community Shannon diversity + season + max_diam

For disease PC2 (Fig. 5, Supp. Table R22), the models were as follows:

M7: disease PC2 ~ log10(bacterial load) + Land use + season + max_diam + (1|Year:Population)

M8: disease PC2 ~ Bacterial Shannon diversity + Land use + season + max_diam + (1|Year:Population)

M9: disease PC2 ~ log10(bacterial load) * Land use + Companion plant community Shannon diversity + season + max_diam + (1|Year:Population)

M10: disease PC2 ~ log10(fungal load) * Land use + season + max_diam + (1|Year:Population)

M11: disease PC1 ~ Fungal Shannon diversity + Land use + season + max_diam + (1|Year:Population)

To assess if disease structured the endophyte microbiome (Supp. Fig. D1), we added disease PC1 into the PERMANOVA using modified Aitchison distance for 16S and ITS data sets as follows. The PERMANOVA was blocked by Population. The significance was assessed by each term in the sequential order of the model below:

Aitchison distance ~ Season + Year + Land use + Soil chemistry PC1 + Companion plant beta-diversity PCoA1 + Mean temperature + Precipitation + log10(reads)

For the Pseudomonas strain level variation (i.e., *HSD* amplicons) associated with disease (Supp. Fig. D2), we used weighted UniFrac distance (with phylogenetic tree rooted at the midpoint) and NMDS ordination. We only included samples with weighted UniFrac distance >= 0.90 and NMDS to  avoid the horseshoe effect and other distributions that challenge the use of

PERMANOVA to partition variance. Weighted UniFrac distance was used here to assess the specific phylogenetic distance within the Pseudomonas genus to best identify strain-level variation that might be associated with disease. As we had fewer samples for which we successfully amplified sufficient *HSD* reads, the model for PERMANOVA was different than the other endophytes and focused specifically on disease:

Weighted UniFrac ~ Season + Land use + disease PC1

We also examined if the presence of HR-like symptoms were associated with differences in community structure for the *HSD* amplicon. The model for the PERMANOVA was as follows:

Weighted UniFrac ~ Season + Land use + HR

<u>Testing for associations between the microbiome and disease symptoms using path analysis</u>
To explore the direct and indirect associations between non-microbial variation (land use type, season, population, year, plant size, companion plant Shannon diversity) and microbial variation (bacteria/fungal Shannon diversity, bacterial/fungal load) on disease, we used structural equation modeling. We split the dataset by season, and Disease PC1 and PC2 were analyzed separately. We used the best predictors from the mixed linear effects modeling to develop the hypotheses tested in models below. Path analysis was performed using Lavaan in R. Bacterial and fungal load were log 10 transformed. Variance was inspected to ensure it was within an order of magnitude; no scaling was necessary. Model diagnostics to assess fit including $X^2$ goodness of fit ($p > 0.05$), CFI (CFI > 0.95), and RMSEA (<0.08). We note that for fall: Disease PC2 ~ fungi; the $X^2$ goodness of fit was marginally significant ($p=0.04$), but this reflects that we cannot predict disease with the measures characterized. The models used are shown below (same between fall and spring, differed by disease PC).

For bacteria and disease PC1, we set residual covariance between bacterial diversity and load to ensure appropriate model fit diagnostics. Analysis was run separately between fall and spring. We fit four models.

1. Disease PC1 ~ Land use + Bacterial Shannon diversity + Bacterial load + Companion plant Shannon diversity + Plant size + Population + Year

2. Bacterial Shannon diversity ~ Land use + Companion plant Shannon diversity + Plant size + Population + Year

3. Bacterial load ~ Land use + Land use + Companion plant Shannon diversity + Plant size + Population + Year

4. Companion plant Shannon diversity ~ Land use + Population + Year

Set residual covariance: Bacterial Shannon diversity ~~ Bacterial load

For fungi and disease PC2, we set residual covariance between fungal diversity and fungal load to ensure appropriate model fit diagnostics. Analyses were run separately for fall and spring. We fit 4 models.

1. Disease PC2 ~ Land use + Fungal Shannon diversity + Fungal load + Companion plant Shannon diversity + Plant size + Population + Year

2. Fungal Shannon diversity ~ Land use + Companion plant Shannon diversity + Plant size + Population + Year

3. Fungal load ~ Land use + Land use + Companion plant Shannon diversity + Plant size + Population + Year

4. Companion plant Shannon diversity ~ Land use + Population + Year

Set residual covariance: Fungal Shannon diversity ~~ Fungal load

**Supp. Table S1:** *matK* primer sequences and PCR cycling conditions

| Sequences_forward | Sequences_reverse |
|---|---|
| CGTACCGTGCTTTTATGTTTACGAG | ATCCTATTCATTTGGAAATCTTGGTGC |
| CGCACAGCGCTTTTGTGTTTACGAG | ACCCCATCCATYTAGAAATCTTGRTTC |
| CGTAYACTRCTTTTATGTTTACGGG | ACCCCGTCCATCTRGAAATYTTRGTTC |
| CGTACAGKACTTTTGTGTTTMCGRG | ATCCCATCCATTTGGAAATYGTGGTTC |
| CGTACAGTMCTTTTGTGTTTACGAG | AYCCCATYCATCTBGAAAAATTGGTYC |
| TGTAKATTRGTTTTGTGTTTATGAG | ATCCTATCCATTTKGAAATCTTGGTTC |
| CGTACAGTACTTTTGTGTTTACGAG | ACCCTGCCCATYTGGAAATHCTAGTTS |
| CGTACAGTACTTTTGTGTTTACAAG | ACCCSATYCATCTMGAAAAAKTGGTTC |
| CGTACACTACTTTTGTGTTTCCGAG | ACCCAGTCCATCTGGAAATCTTGGTTC |
| CGTACSGTGCTTTTATGTTTACGAG | ATCCTATCCATCTGGAAATYTTAGTTC |
| CGTACASTACTTTTGTGTTTACGTG | ACCCTGTYCATCCAGAAATYTTGGTTC |
| CGTACAGTGCTTTTGTGTTTACGAG | ATYCTATMCATTTGGAAATCTTGGTTC |
| CGTACAGTACTTTTATGCTTACGAG | ATCCGATCCATTTKGAACTAWTAGTYC |
| CGTACAGTACTTTTGTGTTTCCKRG | ACCCCATCCATCTAGAAATMGTTRTTC |
| | YTCCYATYCATCTCGARATATTGRTKC |
| | ACCCRGCCCATCTGGAAATCTTGGTTC |
| | ATCCTATCCATCYGGRAATCTTARTTC |
| | ACCCTGTCCATGTGGAAATMTYGATTC |
| | GAACTAAGATWTCCAGATGGATAGGAT |

PCR cycling conditions

| | | |
|---|---|---|
| 95°C | 2min | |
| 95°C | 25sec | x5 |
| 46°C | 35sec | |
| 70°C | 1min | |
| 95°C | 25sec | x35 |
| 48°C | 35sec | |
| 70°C | 1min | |
| 72°C | 5min | |

**Supp. Table S2:** Populations and their corresponding weather stations.

| Population | Weather Station ID |
|---|---|
| Site_01 | KMIHESPE5 |
| Site_02 | KMIHESPE5 |
| Site_03 | KMIHESPE5 |
| Site_04 | KMIWHITE62 |
| Site_05 | KMIMUSKE106 |
| Site_06 | KMIBENTO27 |
| Site_07 | KMIBENTO13 |
| Site_08 | KMIBENTO27 |
| Site_09 | KMIBENTO27 |
| Site_10 | KMISTJOS42 |
| Site_11 | KMISTEVE23 |
| Site_12 | KMISTEVE23 |
| Site_13 | KMITHREE25 |
| Site_14 | KMIHICKO8 |
| Site_15 | KMIHICKO8 |
| Site_16 | KMICOLOM14 |
| Site_17 | KMIBRIDG8 |
| Site_18 | KMIBRIDG23 |
| Site_19 | KMICOLOM14 |

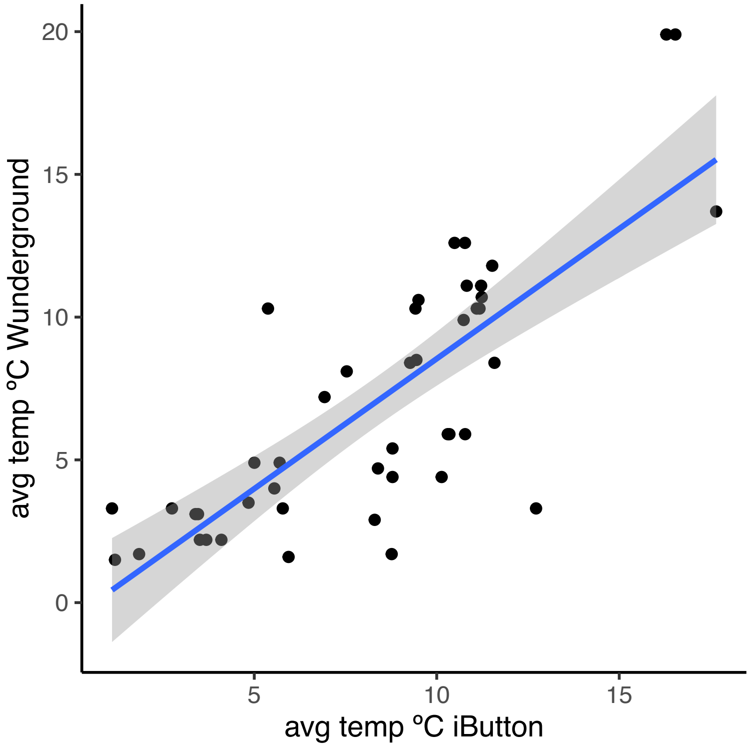


**Supp. Fig. S1:** Temperature was correlated between the iButton measurement and publicly available data from Weather Underground.

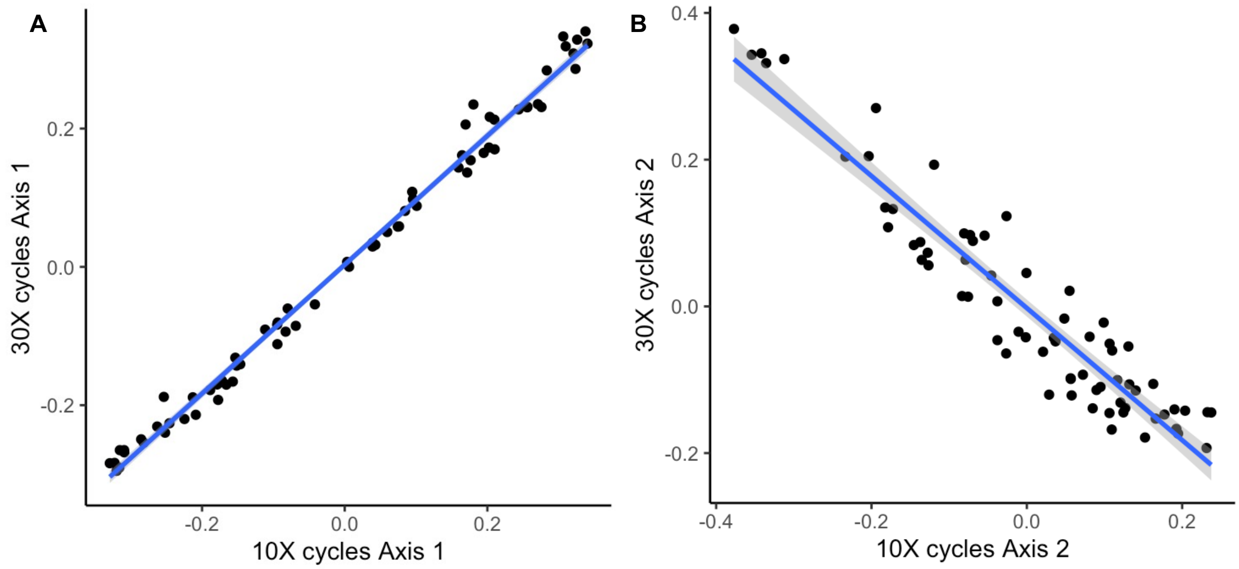


**Supp. Fig. S2:** Cycle number in the first PCR during library preparation does not impact inference of microbiome composition. Points represent values from PCoA Axis 1 (A) and PCoA Axis 2 (B) based on Bray-Curtis dissimilarity from soil samples amplified with either 10 cycles or 30 cycles. Points are well correlated for both Axis 1 (adj. $R^2$ = 0.99, $F_{1,67}$ = 7076, p < 2.2e-16) and Axis 2 (adj. $R^2$ = 0.89, $F_{1,67}$ = 574.9, p < 2.2e-16). Furthermore, using PERMANOVA, cycle only explained less than 1% of variance, while sample ID explained 84.5% of variance.

**Supp. Table M1:** Populations, sample sizes per season, latitude, longitude, and dates of sampling. NA under seasons indicates that apopulation was not sampled.

| Pop. | Y1_Fall | Y1_Spring | Y2_Fall | Y2_Spring | Lat. N | Long. W | Y1_F | Y1_S | Y2_F | Y2_S |
|---|---|---|---|---|---|---|---|---|---|---|
| Site_01 | 16 | 16 | 24 | 24 | 43.52635 | 86.1846 | 11/6/21 | 4/8/22 | 11/8/22 | 4/8/23 |
| Site_02 | 16 | 16 | 24 | 24 | 43.52514 | 86.1843 | 11/6/21 | 4/8/22 | 11/8/22 | 4/8/23 |
| Site_03 | 16 | 16 | 24 | 24 | 43.51874 | 86.17472 | 11/7/21 | 4/8/22 | 11/9/22 | 4/8/23 |
| Site_04 | 16 | 16 | 0 | 24 | 43.34473 | 86.39744 | 11/7/21 | 4/7/22 | NA | 4/7/23 |
| Site_05 | 16 | 16 | 24 | 24 | 43.24858 | 86.33673 | 11/8/21 | 4/7/22 | 11/9/22 | 4/7/23 |
| Site_06 | 16 | 16 | 0 | 24 | 42.09155 | 86.3579 | 11/9/21 | 4/5/22 | NA | 4/2/23 |
| Site_07 | 16 | 16 | 0 | 24 | 42.09834 | 86.32055 | 11/9/21 | 4/5/22 | NA | 4/3/23 |
| Site_08 | 16 | 16 | 24 | 24 | 42.0848 | 86.35274 | 11/9/21 | 4/5/22 | NA | 4/3/23 |
| Site_09 | 16 | 16 | 24 | 24 | 42.0847 | 86.3566 | 11/9/21 | 4/5/22 | 11/10/22 | 4/3/23 |
| Site_10 | 16 | 16 | 24 | 24 | 42.05113 | 86.5091 | 11/10/21 | 4/6/22 | 11/12/22 | 4/4/23 |
| Site_11 | 16 | 16 | 24 | 24 | 42.03627 | 86.51064 | 11/10/21 | 4/3/22 | 11/12/22 | 4/4/23 |
| Site_12 | 16 | 16 | 21 | 24 | 42.03445 | 86.51081 | 11/10/21 | 4/3/22 | 11/12/22 | 4/4/23 |
| Site_13 | 16 | 16 | 23 | 24 | 41.86476 | 86.64401 | 11/10/21 | 4/6/22 | 11/13/22 | 4/4/23 |
| Site_14 | 16 | 16 | 24 | 24 | 42.4031 | 85.39787 | 11/11/21 | 4/4/22 | 11/11/22 | 4/6/23 |
| Site_15 | 16 | 16 | 12 | 23 | 42.40874 | 85.39096 | 11/11/21 | 4/4/22 | 11/11/22 | 4/6/23 |
| Site_16 | 16 | 16 | 0 | 24 | 42.18389 | 86.35902 | 11/11/21 | 4/4/22 | NA | 4/6/23 |
| Site_17 | 16 | 0 | 0 | 0 | 41.87831 | 86.60595 | 11/12/21 | NA | NA | NA |
| Site_18 | 16 | 16 | 24 | 24 | 41.96173 | 86.55059 | 11/12/21 | 4/6/22 | 11/13/22 | 4/5/23 |
| Site_19 | 0 | 16 | 24 | 24 | 42.18366 | 86.38223 | NA | 4/9/22 | 11/11/22 | 4/5/23 |

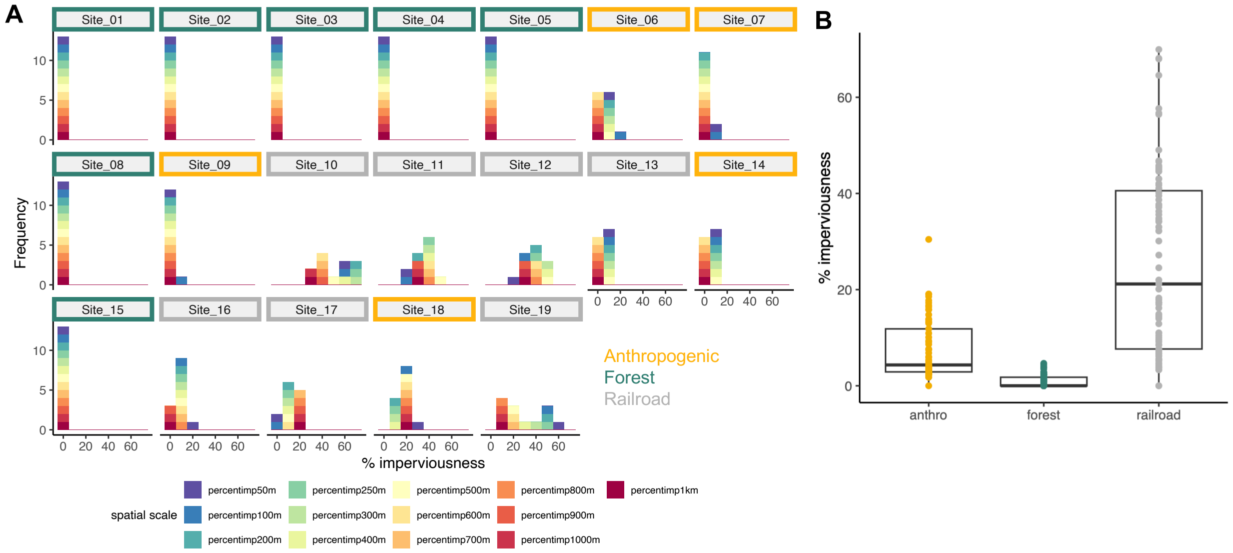


**Supp. Fig. M1**: Imperviousness values to characterize land use types. A) Histogram (bins = 10%) showing the frequency of the percentage imperviousness at different spatial scales, where each color presents the spatial scale. Plots are faceted by site, and the colored outline of the facet label reflects the land use type. B) Railroad populations have highest imperviousness levels at all spatial scales (Kruskall-Wallis $X^2$ = 241.0, df = 25, p = 0.04).

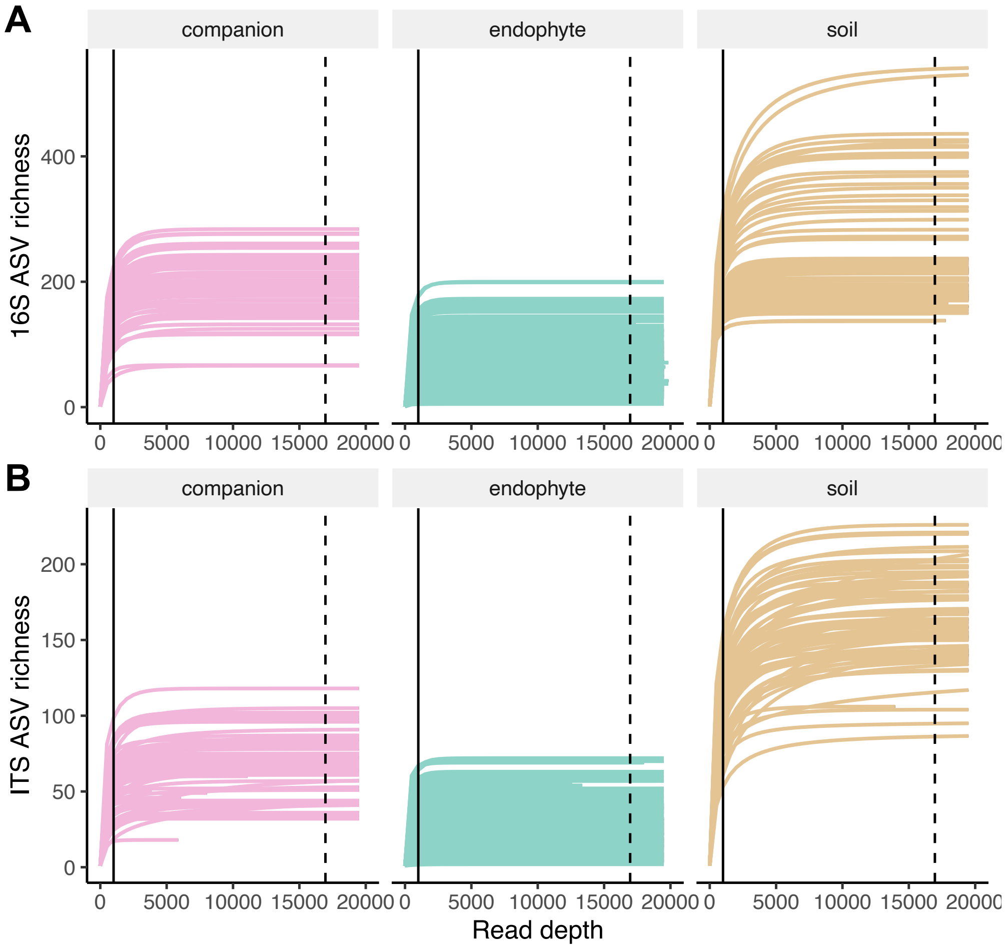


**Supp. Fig. M2:** Rarefaction curves across sample types for A) bacterial and B) fungal microbiomes. The solid line represents the rarefaction depth for endophytes (1000 reads/sample) and dashed lines for companion plants and soil (16963 reads/sample).

**Supp. Table R1**: Fixed effects for *A. thaliana* population size. Significance was evaluated using Type II Wald $X^2$ tests with Kenward-Roger degrees of freedom.

| | Chisq | Df | Pr(>Chisq) |
|---|---|---|---|
| Land use | 1.5791 | 2 | 0.45405 |
| Season | 3.3866 | 1 | 0.06573 |

**Supp. Table R2**: Fixed effects for *A. thaliana* plant size (max. diameter). Significance was evaluated using Type II Wald $X^2$ tests with Kenward-Roger degrees of freedom.

| | Chisq | Df | Pr(>Chisq) |
|---|---|---|---|
| Land use | 1.6176 | 2 | 0.4454 |
| Season | 123.2521 | 1 | <2e-16*** |

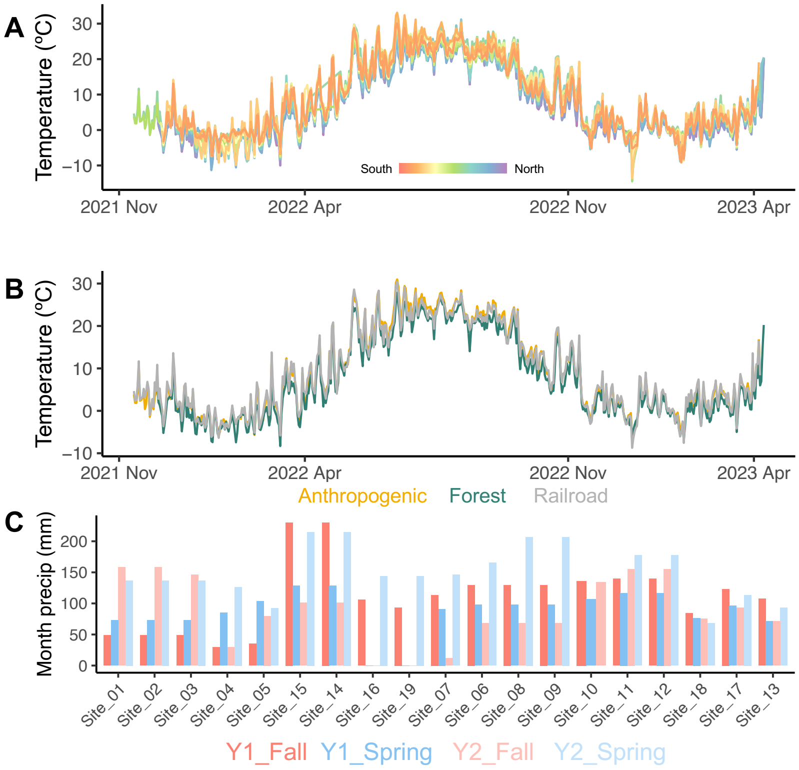


**Supp. Fig. R1:** Climatic variables across sites. A) Temperature profiles from the sampling period. Each line represents a site, colored by latitude. B) The same temperature profiles, but colored by land use type. C) Monthly cumulative precipitation across sites, colored by sampling season.

**Supp. Table R3**: Fixed effects for testing whether temperature and precipitation differ between land use types and/or seasons. Significance was evaluated using Type II Wald $X^2$ tests with Kenward-Roger degrees of freedom.

| | ***Temperature*** | | |
|---|---|---|---|
| | **Chisq** | **Df** | **Pr(>Chisq)** |
| Land use | 1.5759 | 2 | 0.4548 |
| Latitude | 0.5242 | 1 | 0.469 |
| | ***Precipitation*** | | |
| | **Chisq** | **Df** | **Pr(>Chisq)** |
| Land use | 0.3577 | 2 | 0.8362 |
| Season | 27.4812 | 3 | 4.67E-06*** |

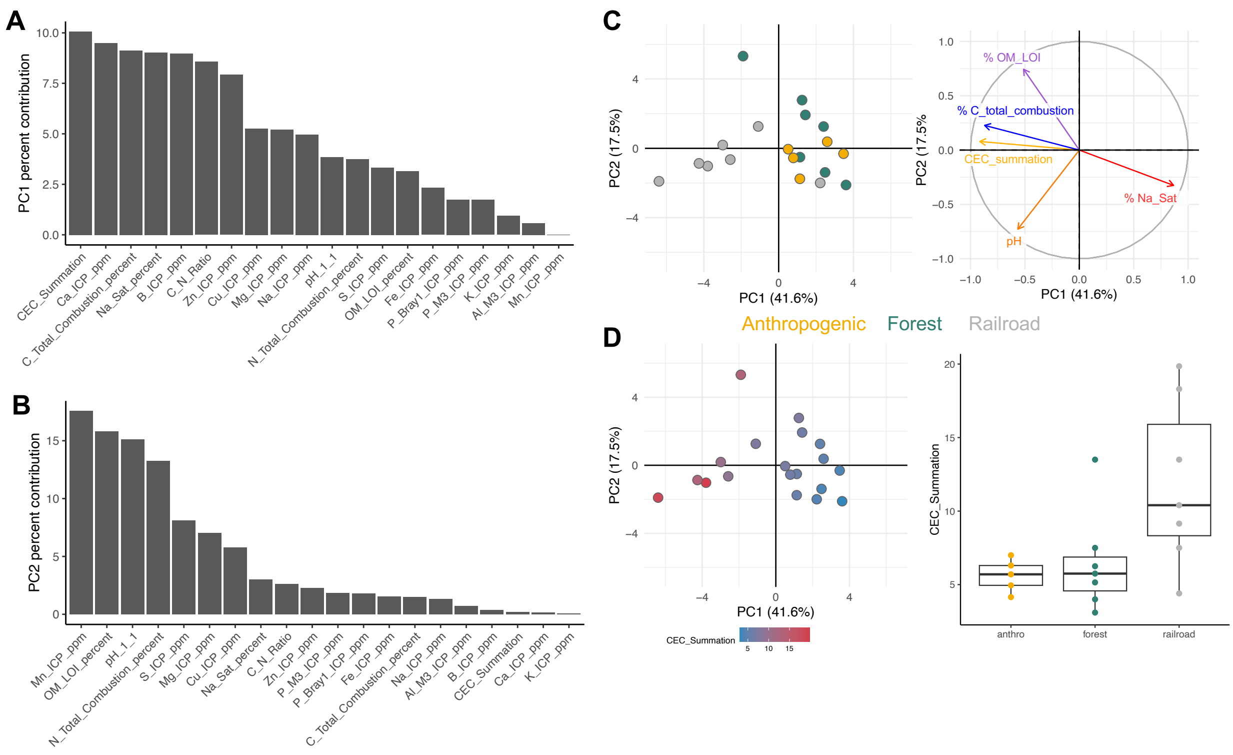


**Supp. Fig. R2:** Principal components analysis for soil chemical properties. A) Percent contribution of different properties to PC1. B) Percent contribution of different properties to PC2. C) PCA biplot showing the sites colored by land use on the right, and the left panel shows the top five most important loadings. D) Cation exchange capacity (CEC_summation), the property that contributed the most to PC1, values displayed in the same PCA plot as in C. CEC_summation was higher in railroad populations compared to the other sample types.

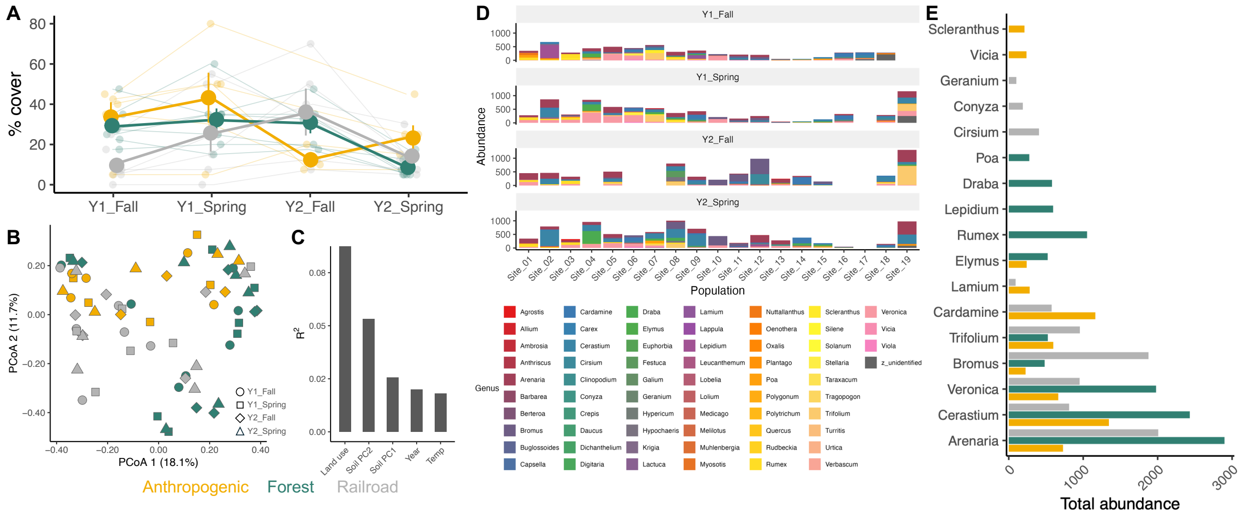


**Supp. Fig. R3:** Companion plant community diversity. A) Percent cover from the companion plant communities. Thin lines and small points show each population, while thick lines and points +/- standard error show the average from each land use category. B) PCoA plot using Bray-Curtis dissimilarity for companion plant community. Colors show land use type, and shapes represent the sampling season. C) $R^2$ values explaining significant variance in companion plant community structure using PERMANOVA. D) Bar plots showing the abundance of the different companion plant genera faceted by sampling season. E) Top 10 most abundant companion plant species across all seasons, colored by land use type.

**Supp. Table R4**: Fixed effects for analysis of percent coverage of the companion plant community. Significance was evaluated using Type II Wald $X^2$ tests with Kenward-Roger degrees of freedom.

| | Chisq | Df | Pr(>Chisq) |
|---|---|---|---|
| Land use | 3.6685 | 2 | 0.1597 |
| Season | 1.1367 | 1 | 0.2864 |
| Soil chemistry PC1 | 1.4457 | 1 | 0.2292 |
| Soil chemistry PC2 | 0.0981 | 1 | 0.7541 |

**Supp. Table R5**: PERMANOVA results for companion plant community beta-diversity based on Bray-Curtis distance, blocked by Population. Terms were evaluated sequentially in the order listed from top to bottom in the table.

| | df | SumofSqs | $R^2$ | F | p value |
|---|---|---|---|---|---|
| Year | 1 | 0.4842 | 0.01995 | 1.5541 | 0.005** |
| Season | 1 | 0.3236 | 0.01333 | 1.0386 | 0.065 |
| Land use | 2 | 2.2541 | 0.09287 | 3.6172 | 0.001*** |
| Soil chemistry PC1 | 1 | 0.6228 | 0.02566 | 1.9988 | 0.001*** |
| Soil chemistry PC2 | 1 | 1.2919 | 0.05323 | 4.1463 | 0.003** |
| Mean temperature | 1 | 0.4364 | 0.01798 | 1.4007 | 0.011* |
| Precipitation | 1 | 0.7869 | 0.03242 | 2.5255 | 0.089 |
| Residual | 58 | 18.0718 | 0.74456 | | |
| Total | 66 | 24.2718 | 1 | | |

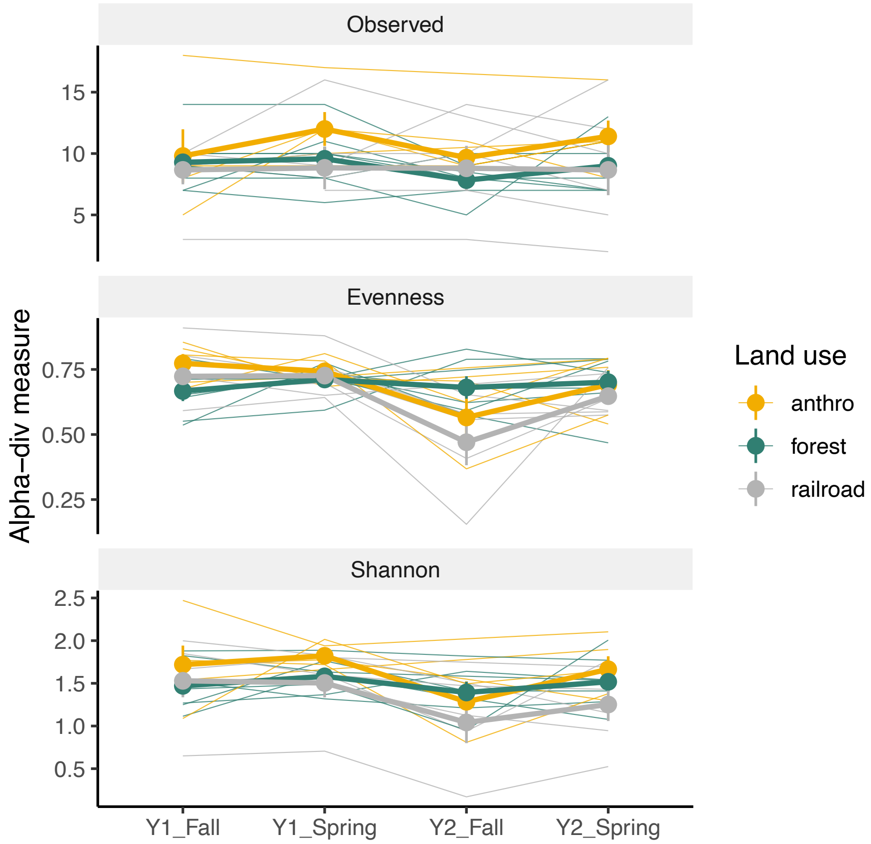


**Supp. Fig. R4:** Alpha-diversity measures for the companion plant community. There was no difference for any measure between seasons or land use types. Thin lines and small points show each population, while thick lines and points +/- standard error show the average from each land use category.

**Supp. Table R6:** Fixed effects for analysis of alpha-diversity measures of the companion plant community. Significance was evaluated using Type II Wald $X^2$ tests with Kenward-Roger degrees of freedom.

| | ***Richness*** | | |
|---|---|---|---|
| | **Chisq** | **Df** | **Pr(>Chisq)** |
| Land use | 3.8807 | 2 | 0.1437 |
| Season | 2.6629 | 1 | 0.1027 |
| | ***Shannon*** | | |
| | **Chisq** | **Df** | **Pr(>Chisq)** |
| Land use | 4.797 | 2 | 0.09085 |
| Season | 2.7274 | 1 | 0.09864 |
| | ***Evenness*** | | |
| | **Chisq** | **Df** | **Pr(>Chisq)** |
| Land use | 1.9331 | 2 | 0.38039 |
| Season | 2.7449 | 1 | 0.09757 |

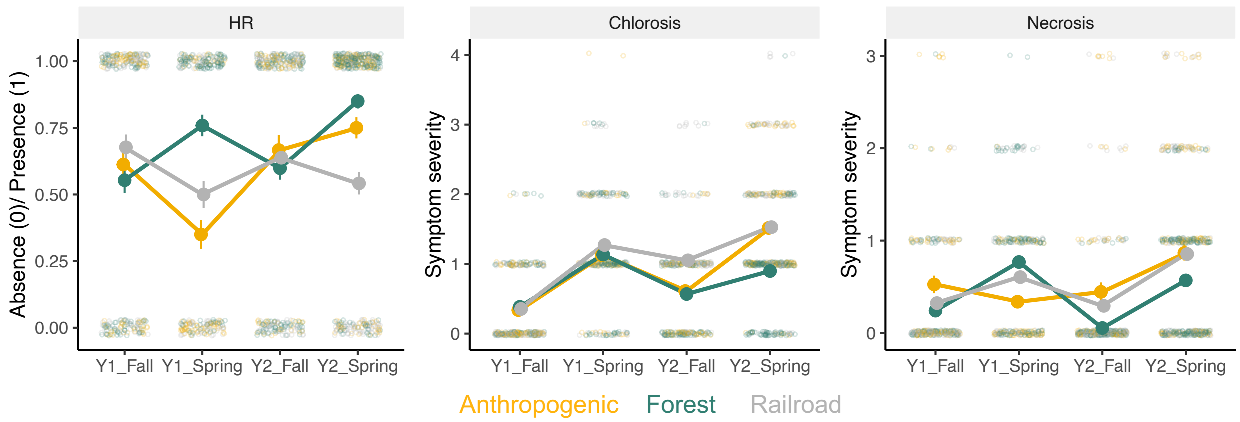


**Supp. Fig. R5:** Seasonal dynamics for the three most commonly observed disease symptoms, hypersensitive response-like lesions (HR), chlorosis, and necrosis. Small points represent individual plants. Lines and points +/- standard error show the average from each land use category.

**Supp. Table R7A:** Fixed effects for analysis of disease symptoms over land use and season, and companion plant community diversity. Significance was evaluated using Type III Wald $X^2$ tests with Kenward-Roger degrees of freedom.

| | Chisq | Df | Pr(>Chisq) |
|---|---|---|---|
| (Intercept) | 0.1973 | 1 | 0.6569 |
| Land use | 0.1093 | 2 | 0.9468 |
| Season | 32.4282 | 1 | 1.24E-08*** |
| Comp. plant Shannon | 0.3525 | 1 | 0.5527 |
| Plant size (max diam) | 25.339 | 1 | 4.81E-07*** |
| Land use * Season | 35.7356 | 2 | 1.74E-08*** |
| Land use * Comp. plant Shannon | 0.1985 | 2 | 0.9055 |
| Season * Comp. plant Shannon | 24.073 | 1 | 9.28E-07*** |
| Land use * Comp. plant Shannon * Season | 33.3112 | 2 | 5.84E-08*** |

**Supp. Table R7B:** Model selection for disease symptom dynamics.

Model1: Disease PC1 ~ Land use * Season + Plant size + (1|Year:Population)

Model2: Disease PC1 ~ Land use * Season * Comp. plant Shannon + Plant size + (1|Year:Population)

| | npar | AIC | BIC | logLik | trans log lik | Chisq | Df | Pr(>Chisq) |
|---|---|---|---|---|---|---|---|---|
| Model1 | 9 | 1914.1 | 1960.8 | -948.03 | 1896.1 | | | |
| Model2 | 15 | 1887.2 | 1965.1 | -928.62 | 1857.2 | 38.836 | 6 | 7.71E-07*** |

**Supp. Table R8:** Fixed effects for analysis of microbial load. Significance was evaluated using Type II Wald $X^2$ tests with Kenward-Roger degrees of freedom.

| | | | |
|---|---|---|---|
| | ***Bacteria*** | | |
| | **Chisq** | **Df** | **Pr(>Chisq)** |
| Season | 463.5314 | 1 | < 2.20E-16*** |
| Land use | 0.1863 | 2 | 0.911076 |
| Plant size | 8.0195 | 1 | 0.004628** |
| | ***Fungi*** | | |
| | **Chisq** | **Df** | **Pr(>Chisq)** |
| Season | 5.9241 | 1 | 0.01494* |
| Land use | 0.7137 | 2 | 0.69987 |
| Plant size | 1.2773 | 1 | 0.2584 |

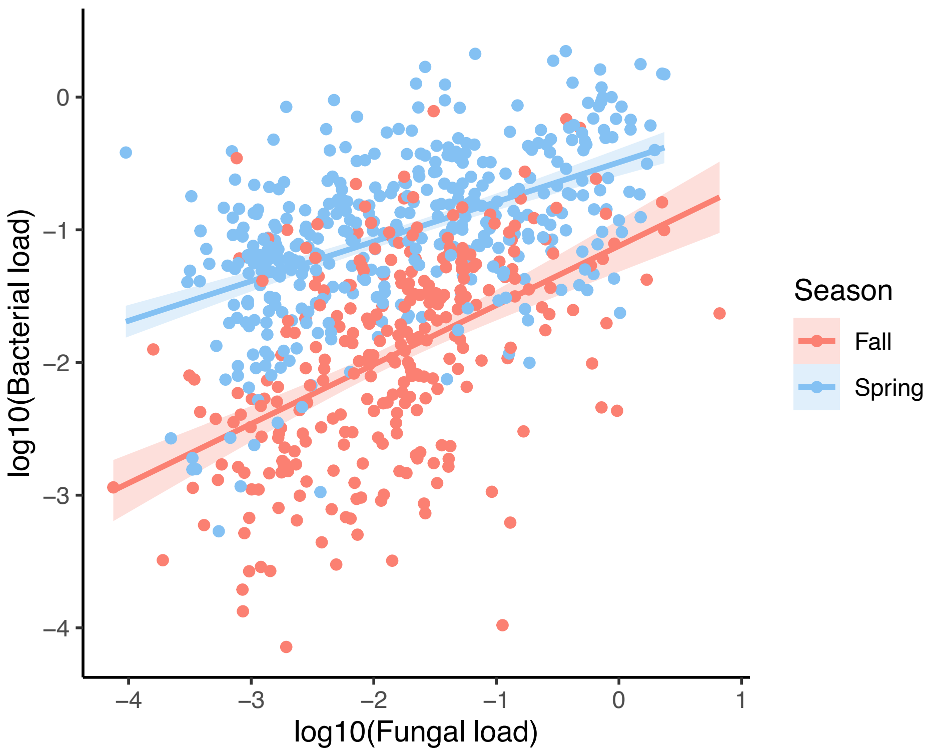


**Supp. Fig. R6:** Bacterial and fungal loads are positively correlated, and the strength of this correlation differed between fall and spring. There was no effect of land use on this correlation. Each point represents an individual *A. thaliana* endophyte, colored by season.

**Supp. Table R9:** Fixed effects for the analysis of the correlation between bacterial and fungal load. Significance was evaluated using Type II Wald $X^2$ tests with Kenward-Roger degrees of freedom.

| | **Chisq** | **Df** | **Pr(>Chisq)** |
|---|---|---|---|
| log10(fungal load) | 60.1101 | 1 | 8.97E-15*** |
| Season | 490.4194 | 1 | < 2.20E-16*** |
| Plant size | 1.7237 | 1 | 0.1892 |
| log10(fungal load) * Season | 23.8971 | 1 | 1.02E-06*** |

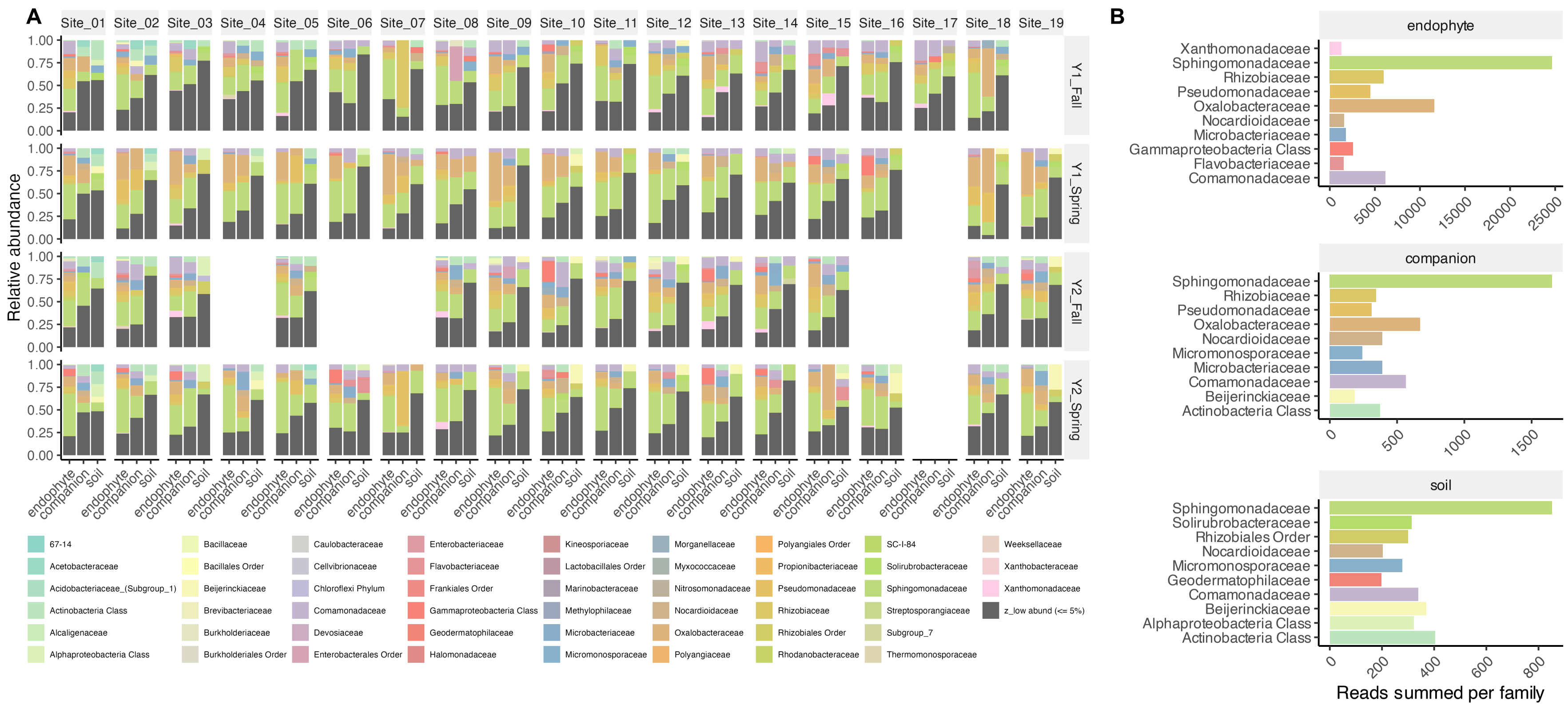


**Supp. Fig. R7:** Bacterial family relative abundance. A) Barplots show the relative abundance by the family level for endophytes, companion plants, and soil. Plots are faceted by site and season. Dark grey bars represent all ASVs at <5% relative abundance. B) The top 10 most abundant bacterial families for each sample type across all season. Colors in A and B match.

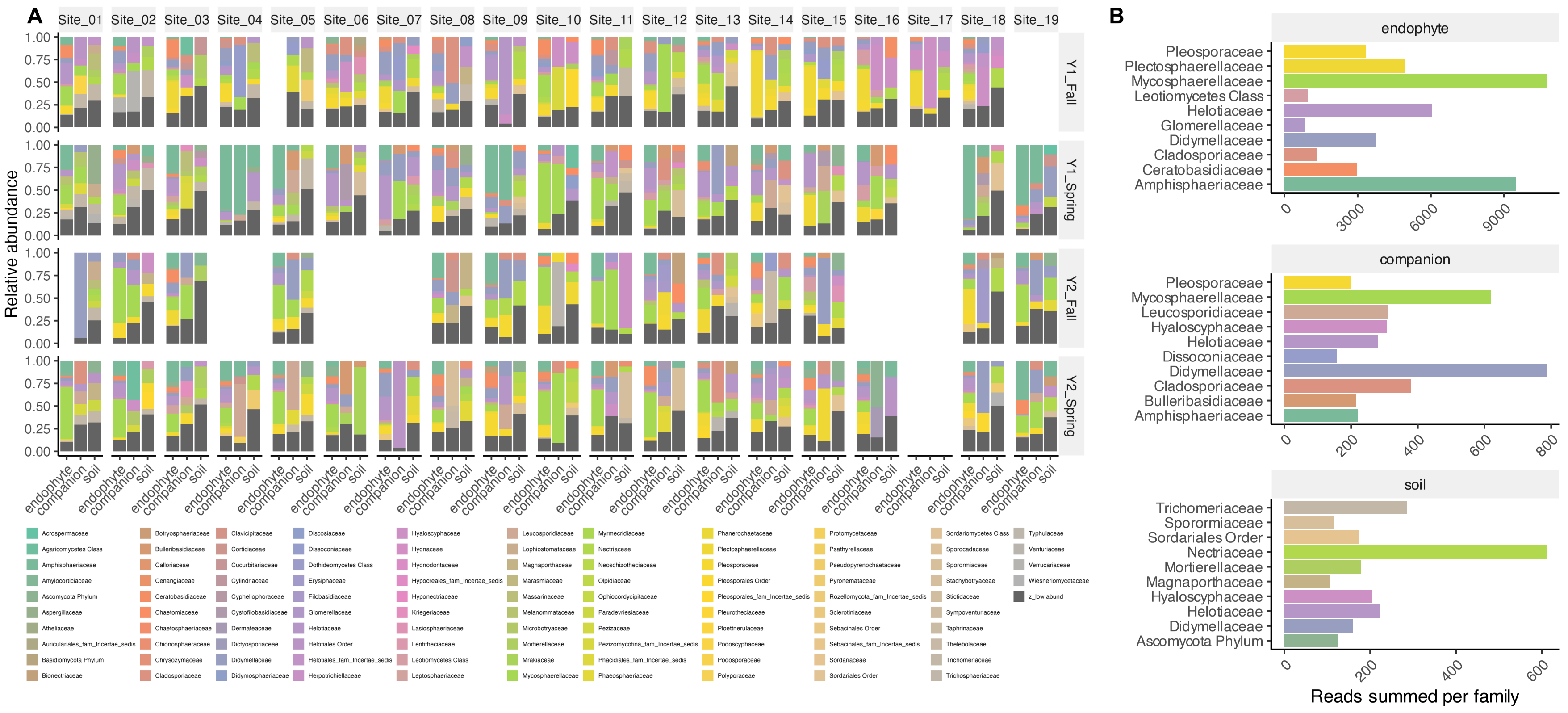


**Supp. Fig. R8**: Fungal family relative abundance. A) Barplots show the relative abundance by the family level for endophytes, companion plants, and soil. Plots are faceted by site and season. Dark grey bars represent all ASVs at <5% relative abundance. B) The top 10 most abundant bacterial families for each sample type across all season. Colors in A and B match.

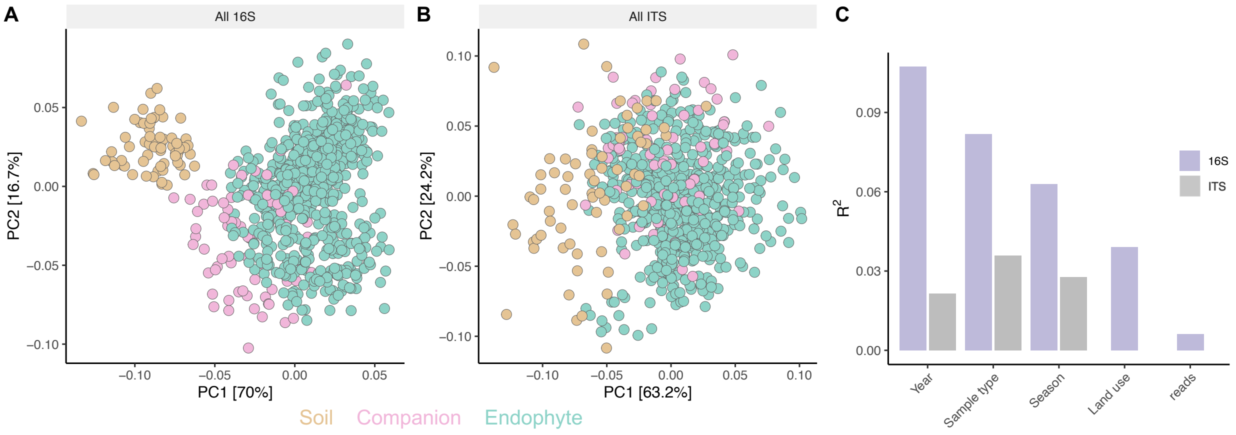


**Supp. Fig. R9:** Beta-diversity PCoA plots for all sample types using modified Aitchinson distance matrix. A) Bacterial microbiomes. B) Fungal microbiomes. For A-B, each green point represents an individual A. thaliana endophyte, while the companion plants (blue) and soil (brown) are pools per site and season. C) PERMANOVA results with the percent variance ($R^2$) for terms in the model.

**Supp. Table R10:** PERMANOVA results for microbiome beta-diversity using modified Aitchison distance between sample types, blocked by Population. Terms were evaluated sequentially in the order listed from top to bottom in the table.

| | | | ***Bacteria*** | | |
|---|---|---|---|---|---|
| | **df** | **SumOfSqs** | **$R^2$** | **F** | **p value** |
| Sample type | 2 | 159.9 | 0.08178 | 50.7301 | 0.001*** |
| Season | 1 | 122.86 | 0.06283 | 77.9569 | 0.001*** |
| Year | 1 | 209.77 | 0.10729 | 133.1092 | 0.001*** |
| Land use | 2 | 76.31 | 0.03903 | 24.2098 | 0.001*** |
| log10(reads) | 1 | 12.17 | 0.00623 | 7.7232 | 0.002** |
| Residual | 872 | 1374.24 | 0.70285 | | |
| Total | 879 | 1955.24 | 1 | | |
| | | | ***Fungi*** | | |
| | **df** | **SumOfSqs** | **$R^2$** | **F** | **Pr(>F)** |
| Sample type | 2 | 69.65 | 0.03591 | 14.8315 | 0.001*** |
| Season | 1 | 53.86 | 0.02777 | 22.9393 | 0.001*** |
| Year | 1 | 41.56 | 0.02143 | 17.6988 | 0.001*** |
| Land use | 2 | 5.33 | 0.00275 | 1.1353 | 1 |
| log10(reads) | 1 | 0.95 | 0.00049 | 0.4036 | 0.735 |
| Residual | 753 | 1768.11 | 0.91165 | | |
| Total | 760 | 1939.46 | 1 | | |

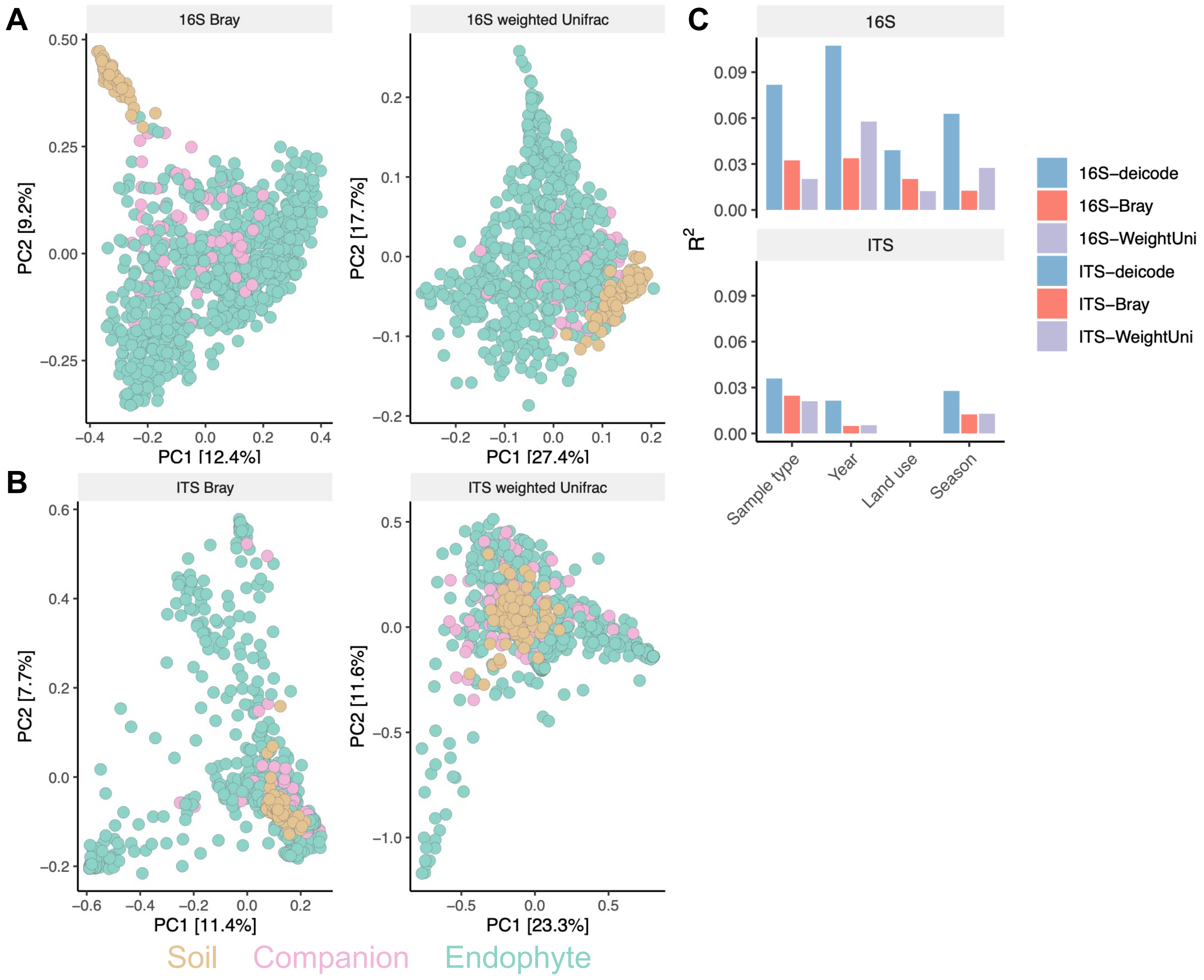


**Supp. Fig. R10:** Ordination analyses for beta-diversity analyses between sample types using different distance metrics show similar patterns to the modified Aitchinson distance metric. A) Beta-diversity for bacterial microbiomes using Bray-Curtis and weighted UniFrac metrics. Points are colored by sample type. B) Beta-diversity for fungal microbiomes using Bray-Curtis and weighted UniFrac metrics. Points are colored by sample type. C) PERMANOVA results with the percent variance ($R^2$) for terms in the model, where the bars represent the distance metric. Blue is the modified Aitchinson distance metric, implement in package DEICODE.

**Supp. Table R11**: PERMANOVA results for microbiome beta-diversity using different distance metrics between sample types, blocked by Population. Terms were evaluated sequentially in the order listed from top to bottom in the table.

| | | | | | |
|---|---|---|---|---|---|
| | | ***Bacteria: Bray-Curtis*** | | | |
| | **Df** | **SumOfSqs** | **$R^2$** | **F** | **Pr(>F)** |
| Sample type | 2 | 7.841 | 0.03237 | 15.1612 | 0.001*** |
| Season | 1 | 3.039 | 0.01255 | 11.7507 | 0.001*** |
| Year | 1 | 8.174 | 0.03375 | 31.6094 | 0.001*** |
| Land use | 2 | 4.896 | 0.02021 | 9.4662 | 0.001*** |
| Residual | 844 | 218.243 | 0.90112 | | |
| Total | 850 | 242.192 | 1 | | |
| | | ***Bacteria: Weighted UniFrac*** | | | |
| | **Df** | **SumOfSqs** | **$R^2$** | **F** | **Pr(>F)** |
| Sample type | 2 | 7.841 | 0.03237 | 15.1612 | 0.001*** |
| Season | 1 | 3.039 | 0.01255 | 11.7507 | 0.001*** |
| Year | 1 | 8.174 | 0.03375 | 31.6094 | 0.001*** |
| Land use | 2 | 4.896 | 0.02021 | 9.4662 | 0.001*** |
| Residual | 844 | 218.243 | 0.90112 | | |
| Total | 850 | 242.192 | 1 | | |
| | | ***Fungi: Bray-Curtis*** | | | |
| | **Df** | **SumOfSqs** | **$R^2$** | **F** | **Pr(>F)** |
| Sample type | 2 | 6.593 | 0.0243 | 8.1684 | 0.001*** |
| Season | 1 | 3.338 | 0.01231 | 8.2719 | 0.001*** |
| Year | 1 | 1.338 | 0.00493 | 3.3154 | 0.001*** |
| Land use | 2 | 0.929 | 0.00343 | 1.1512 | 1 |
| Residual | 642 | 259.082 | 0.95504 | | |
| Total | 648 | 271.281 | 1 | | |
| | | ***Fungi: Weighted UniFrac*** | | | |
| | **Df** | **SumOfSqs** | **$R^2$** | **F** | **Pr(>F)** |
| Sample type | 2 | 5.768 | 0.02045 | 6.8527 | 0.001*** |
| Season | 1 | 3.606 | 0.01278 | 8.5688 | 0.001*** |
| Year | 1 | 1.529 | 0.00542 | 3.6335 | 0.002** |
| Land use | 2 | 0.985 | 0.00349 | 1.1698 | 1 |
| Residual | 642 | 270.18 | 0.95786 | | |
| Total | 648 | 282.068 | 1 | | |

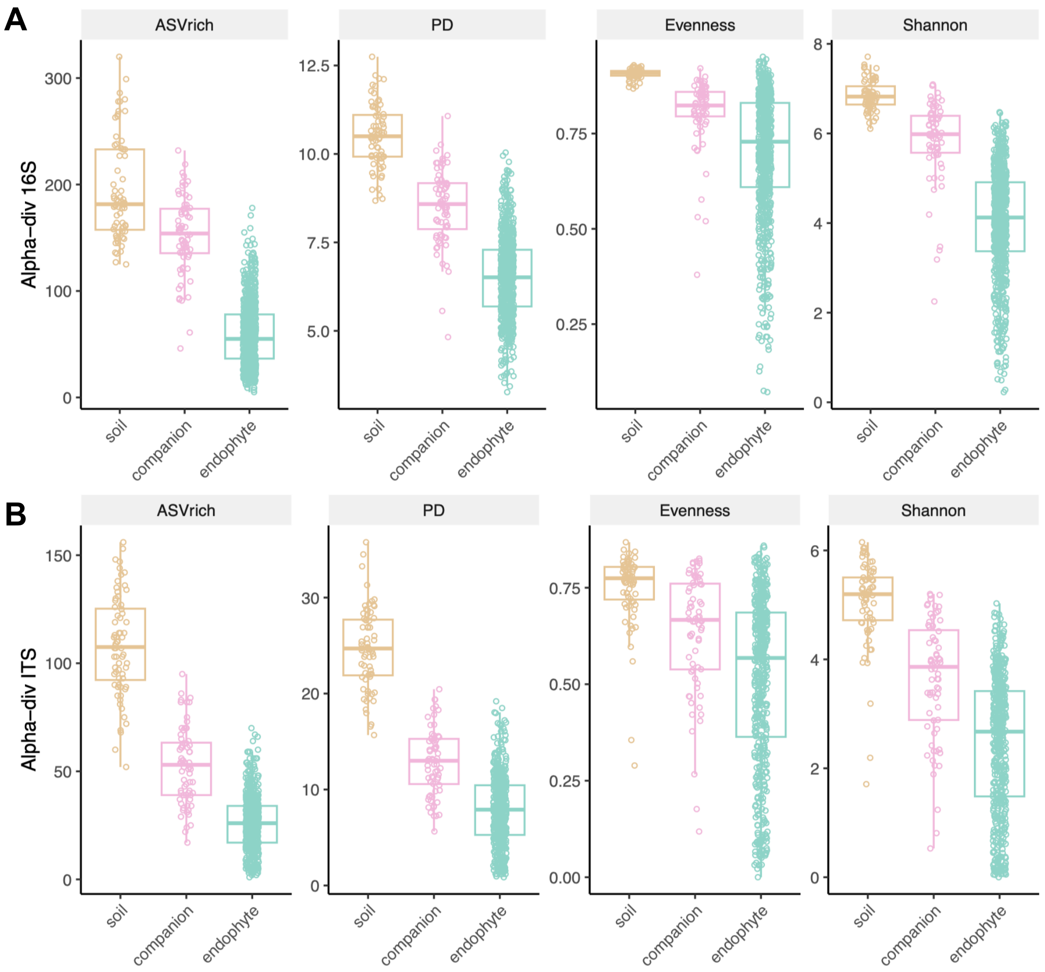


**Supp. Fig. R11:** Alpha-diversity metrics for microbiome diversity for A) bacteria and B) fungi. Each point represents an individual sample. Soil and companion plant samples were pools per site and season. Colors represent sample types as labeled on the x-axis.

**Supp. Table R12:** Alpha-diversity analyses across sample types. All tests used Kruskal-Wallis $X^2$ tests.

| Alpha-div. measure | Kruskal-Wallis $X^2$ | df | p-value |
|---|---|---|---|
| 16S ASV rich | 319.22 | 2 | <0.0001*** |
| 16S Faith's PD | 282.42 | 2 | <0.0001*** |
| 16S Shannon | 287.37 | 2 | <0.0001*** |
| 16S Evenness | 197.01 | 2 | <0.0001*** |
| ITS ASV rich | 266.85 | 2 | <0.0001*** |
| ITS Faith's PD | 247.27 | 2 | <0.0001*** |
| ITS Shannon | 194.5 | 2 | <0.0001*** |
| ITS Evenness | 110.86 | 2 | <0.0001*** |

**Supp. Table R13:** PERMANOVA results for bacterial microbiome beta-diversity using different distance metrics within each sample type, blocked by Population. Terms were evaluated sequentially in the order listed from top to bottom in the table.

| ***16S: A. thaliana endophytes*** | | | | | |
|---|---|---|---|---|---|
| | **Df** | **SumOfSqs** | **$R^2$** | **F** | **Pr(>F)** |
| Season | 1 | 439.02 | 0.27741 | 349.3073 | 0.001*** |
| Year | 1 | 120.13 | 0.07591 | 95.5838 | 0.001*** |
| Land use | 2 | 13.96 | 0.00882 | 5.5534 | 0.001*** |
| Soil chem PC1 | 1 | 32.28 | 0.02039 | 25.6809 | 0.001*** |
| Comp. plant Bray-Curtis PC1 | 1 | 7.83 | 0.00495 | 6.2306 | 0.001*** |
| Mean Temperature | 1 | 22 | 0.0139 | 17.5076 | 0.001*** |
| Precipitation | 1 | 7.77 | 0.00491 | 6.1833 | 0.001*** |
| log10(reads) | 1 | 17.09 | 0.0108 | 13.5956 | 0.001*** |
| Residual | 734 | 922.51 | 0.58291 | | |
| Total | 743 | 1582.58 | 1 | | |
| ***16S: Companion plant microbiome*** | | | | | |
| | **Df** | **SumOfSqs** | **$R^2$** | **F** | **Pr(>F)** |
| Season | 1 | 44.628 | 0.32104 | 36.5141 | 0.001*** |
| Year | 1 | 3 | 0.02158 | 2.4545 | 0.09 |
| Land use | 2 | 11.009 | 0.07919 | 4.5036 | 0.001*** |
| Soil chem PC1 | 1 | 0.353 | 0.00254 | 0.2886 | 0.001*** |
| Comp. plant Bray-Curtis PC1 | 1 | 0.698 | 0.00502 | 0.5714 | 0.746 |
| Mean Temperature | 1 | 3.656 | 0.0263 | 2.9912 | 0.077 |
| Precipitation | 1 | 4.245 | 0.03054 | 3.473 | 0.057 |
| log10(reads) | 1 | 0.534 | 0.00384 | 0.4369 | 0.866 |
| Residual | 58 | 70.889 | 0.50995 | | |
| Total | 67 | 139.011 | 1 | | |
| ***16S: Soil microbiome*** | | | | | |
| | **Df** | **SumOfSqs** | **$R^2$** | **F** | **Pr(>F)** |
| Season | 1 | 1.277 | 0.00907 | 1.9627 | 0.036* |
| Year | 1 | 0.986 | 0.007 | 1.5151 | 0.169 |
| Land use | 2 | 62.176 | 0.44148 | 47.7865 | 0.001*** |
| Soil chem PC1 | 1 | 3.19 | 0.02265 | 4.9031 | 0.001*** |
| Comp. plant Bray-Curtis PC1 | 1 | 19.948 | 0.14164 | 30.6622 | 0.001*** |
| Mean Temperature | 1 | 2.082 | 0.01479 | 3.201 | 0.249 |
| Precipitation | 1 | 1.127 | 0.008 | 1.732 | 0.061 |
| log10(reads) | 1 | 12.319 | 0.08747 | 18.9357 | 0.001*** |
| Residual | 58 | 37.733 | 0.26792 | | |
| Total | 67 | 140.837 | 1 | | |

**Supp. Table R14:** PERMANOVA results for fungal microbiome beta-diversity using different distance metrics within each sample type, blocked by Population. Terms were evaluated sequentially in the order listed from top to bottom in the table.

| ***ITS: A. thaliana endophytes*** | | | | | |
|---|---|---|---|---|---|
| | **Df** | **SumOfSqs** | **$R^2$** | **F** | **Pr(>F)** |
| Season | 1 | 30.73 | 0.02005 | 13.2665 | 0.001*** |
| Year | 1 | 24.86 | 0.01622 | 10.7311 | 0.001*** |
| Land use | 2 | 14.78 | 0.00964 | 3.1911 | 0.319 |
| Soil chem PC1 | 1 | 4.07 | 0.00265 | 1.7551 | 0.234 |
| Comp. plant Bray-Curtis PC1 | 1 | 22.09 | 0.01441 | 9.5381 | 0.001*** |
| Mean Temperature | 1 | 6.42 | 0.00419 | 2.7701 | 0.019* |
| Precipitation | 1 | 3.03 | 0.00197 | 1.3062 | 0.347 |
| log10(reads) | 1 | 2.41 | 0.00157 | 1.0389 | 0.375 |
| Residual | 615 | 1424.53 | 0.9293 | | |
| Total | 624 | 1532.9 | 1 | | |
| ***ITS: Companion plant microbiome*** | | | | | |
| | **Df** | **SumOfSqs** | **$R^2$** | **F** | **Pr(>F)** |
| Season | 1 | 44.438 | 0.24417 | 26.7383 | 0.001*** |
| Year | 1 | 1.025 | 0.00563 | 0.617 | 0.572 |
| Land use | 2 | 24.58 | 0.13505 | 7.3947 | 0.001*** |
| Soil chem PC1 | 1 | 1.038 | 0.0057 | 0.6245 | 0.006** |
| Comp. plant Bray-Curtis PC1 | 1 | 3.724 | 0.02046 | 2.2404 | 0.079 |
| Mean Temperature | 1 | 2.746 | 0.01509 | 1.6521 | 0.159 |
| Precipitation | 1 | 6.453 | 0.03546 | 3.8828 | 0.008** |
| log10(reads) | 1 | 1.603 | 0.00881 | 0.9643 | 0.351 |
| Residual | 58 | 96.394 | 0.52964 | | |
| Total | 67 | 182 | 1 | | |
| ***ITS: Soil microbiome*** | | | | | |
| | **Df** | **SumOfSqs** | **$R^2$** | **F** | **Pr(>F)** |
| Season | 1 | 0.949 | 0.00617 | 0.6266 | 0.282 |
| Year | 1 | 0.899 | 0.00584 | 0.5934 | 0.351 |
| Land use | 2 | 35.158 | 0.22851 | 11.6038 | 0.076 |
| Soil chem PC1 | 1 | 12.403 | 0.08061 | 8.187 | 0.109 |
| Comp. plant Bray-Curtis PC1 | 1 | 8.612 | 0.05598 | 5.6849 | 0.392 |
| Mean Temperature | 1 | 2.27 | 0.01476 | 1.4986 | 0.287 |
| Precipitation | 1 | 1.756 | 0.01141 | 1.1593 | 0.252 |
| log10(reads) | 1 | 3.943 | 0.02563 | 2.6026 | 0.095 |
| Residual | 58 | 87.867 | 0.57109 | | |
| Total | 67 | 153.857 | 1 | | |

**Supp. Table R15:** Fixed effects for the analysis of Shannon diversity in amplicons across different phylogenetic levels for *A. thaliana* endophytes. Significance was evaluated using Type III Wald $X^2$ tests with Kenward-Roger degrees of freedom.

| | **Chisq** | **Df** | **Pr(>Chisq)** |
|---|---|---|---|
| (Intercept) | 623.2662 | 1 | < 2.20E-16*** |
| Amplicon | 215.134 | 3 | < 2.20E-16*** |
| Season | 0.1085 | 1 | 0.7418365 |
| Land use | 2.6099 | 2 | 0.2711909 |
| Plant size | 10.0486 | 1 | 0.0015246** |
| Amplicon * Season | 15.6899 | 3 | 0.0013127** |
| Amplicon * Land use | 16.7126 | 6 | 0.0103995* |
| Season * Land use | 15.4901 | 2 | 0.0004329*** |
| Amplicon * Season * Land use | 15.096 | 6 | 0.0195233* |

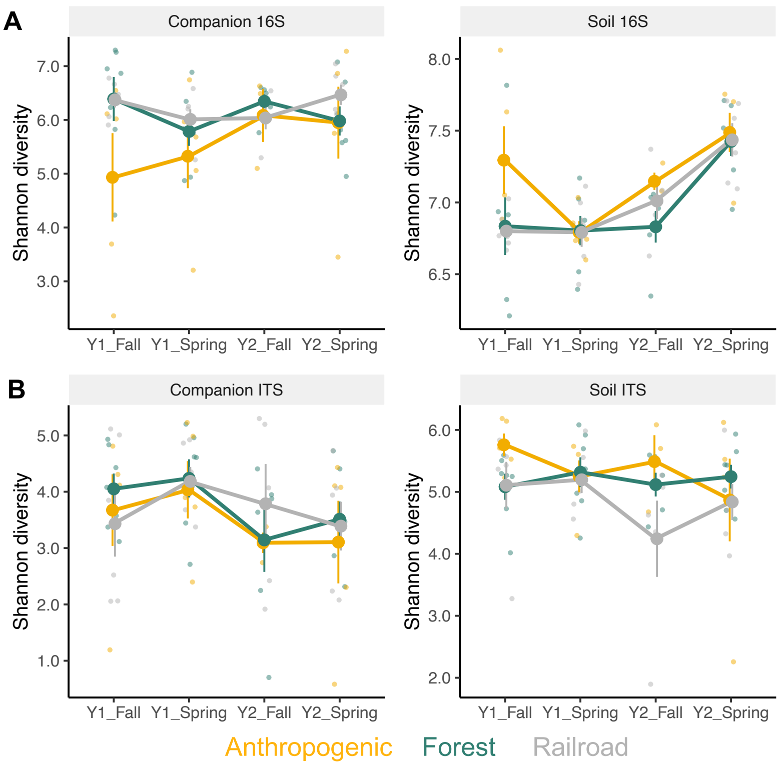


**Supp. Fig. R12:** Temporal dynamics for Shannon diversity in the companion plant and soil microbiomes for bacteria (A) and fungi (B). Small points represent each site measure, while Lines and points +/- standard error show the average from each land use category. Color represents land use.

**Supp. Table R16:** Fixed effects for the analysis of companion plant and soil microbiome diversity across land use and seasons. Significance was evaluated using Type II Wald $X^2$ tests with Kenward-Roger degrees of freedom.

| | | | |
|---|---|---|---|
| ***16S: Companion plant microbiomes*** | | | |
| | **Chisq** | **Df** | **Pr(>Chisq)** |
| Season | 0.1999 | 1 | 0.6548 |
| Land use | 5.779 | 2 | 0.0556 |
| ***16S: Soil microbiomes*** | | | |
| | **Chisq** | **Df** | **Pr(>Chisq)** |
| Season | 2.9032 | 1 | 0.0884 |
| Land use | 2.7298 | 2 | 0.2554 |
| ***ITS: Companion plant microbiomes*** | | | |
| | **Chisq** | **Df** | **Pr(>Chisq)** |
| Season | 0.7621 | 1 | 0.3827 |
| Land use | 0.5141 | 2 | 0.7733 |
| ***ITS: Soil microbiomes*** | | | |
| | **Chisq** | **Df** | **Pr(>Chisq)** |
| Season | 0.1903 | 1 | 0.6626 |
| Land use | 2.1946 | 2 | 0.3338 |

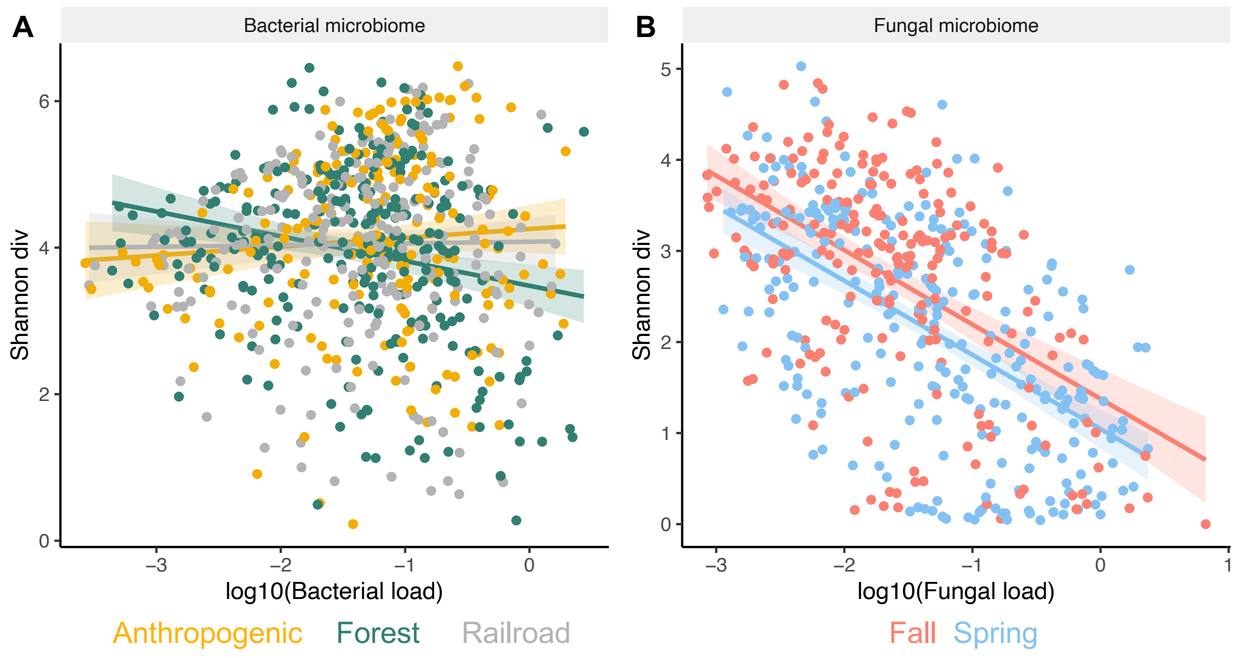


**Supp. Fig. R13:** Correlations between microbial load and diversity. A) Bacterial Shannon diversity and load correlations depended on land use type, but with a negative association the forest, but marginal positive correlations in anthropogenic and railroad populations. B) Fungal Shannon diversity and load were negatively correlated, and this correlation was sensitive to season, but not land use. Points are individual *A. thaliana* endophytes, colored by the factor that explained significant variance.

**Supp. Table R17:** Fixed effects for the analysis of correlations between microbial load and microbial diversity. Significance was evaluated using Type II Wald $X^2$ tests with Kenward-Roger degrees of freedom.

| | ***16S: Endophyte*** | | |
|---|---|---|---|
| | **Chisq** | **Df** | **Pr(>Chisq)** |
| log10(bacterial load) | 1.3583 | 1 | 0.243833 |
| Land use | 1.2764 | 2 | 0.528233 |
| Season | 0.0275 | 1 | 0.86825 |
| Plant size | 0.7816 | 1 | 0.376652 |
| log10(bacterial load) * Land use | 13.0416 | 2 | 0.001472** |
| | ***ITS: Endophyte*** | | |
| | **Chisq** | **Df** | **Pr(>Chisq)** |
| log10(fungal load) | 321.3981 | 1 | < 2.20E-16*** |
| Land use | 0.6969 | 2 | 0.7057876 |
| Season | 11.9104 | 1 | 0.0005582*** |
| Plant size | 0.1952 | 1 | 0.6586307 |

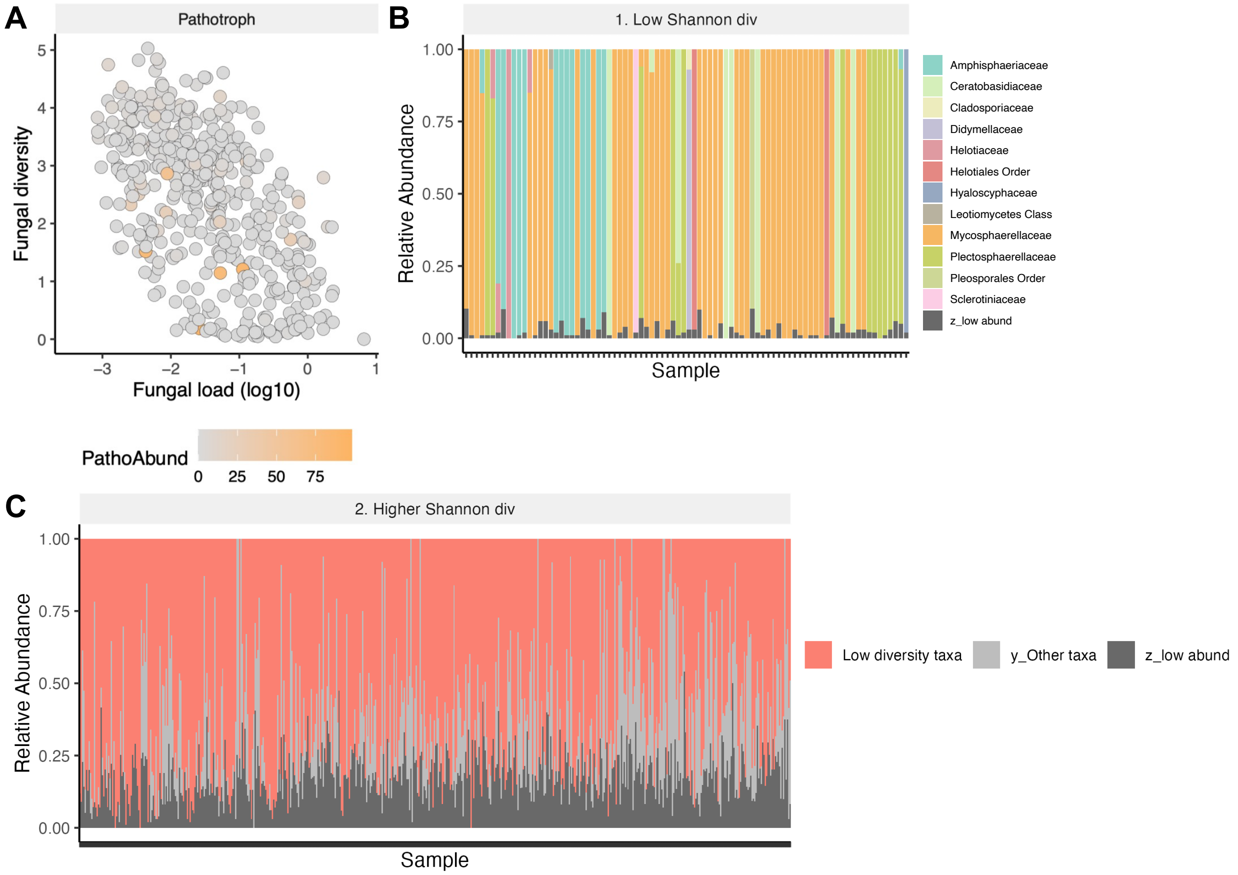


**Supp. Fig. R14:** Low diversity and high fungal load is not associated with trophic mode or taxonomy. A) There is no association between pathotrophs and the negative covariation between fungal load and fungal diversity. Each point is an individual Arabidopsis sample, colored by the relative abundance of pathotrophs. B) 12 different fungal families are associated with low Shannon diversity (<= 1 Shannon diversity). C) However, these fungal families are found across all other samples with higher Shannon diversity, suggesting that pathotrophs or particular taxa are not associated with the negative covariation between Shannon diversity and fungal load.

**Supp. Table R18:** Fixed effects for the analysis of correlations between *A. thaliana* endophyte microbial Shannon diversity and companion plant community Shannon diversity. Significance was evaluated using Type II Wald $X^2$ tests with Kenward-Roger degrees of freedom.

| ***16S: Endophyte microbiome*** | | | |
|---|---|---|---|
| | **Chisq** | **Df** | **Pr(>Chisq)** |
| Comp. plant diversity | 16.7972 | 1 | 4.16E-05*** |
| Land use | 1.8505 | 2 | 0.39642 |
| Season | 3.3895 | 1 | 0.06561 |
| Plant size | 0.6827 | 1 | 0.40866 |
| Comp. plant diversity * Land use | 8.5534 | 2 | 0.01389* |
| ***ITS: Endophyte microbiome*** | | | |
| | **Chisq** | **Df** | **Pr(>Chisq)** |
| Comp. plant diversity | 9.1759 | 1 | 0.002452** |
| Land use | 4.9373 | 2 | 0.0847 |
| Season | 45.3816 | 1 | 1.62E-11*** |
| Plant size | 0.0462 | 1 | 0.8299 |
| Comp. plant diversity * Land use | 6.4895 | 2 | 0.038978* |

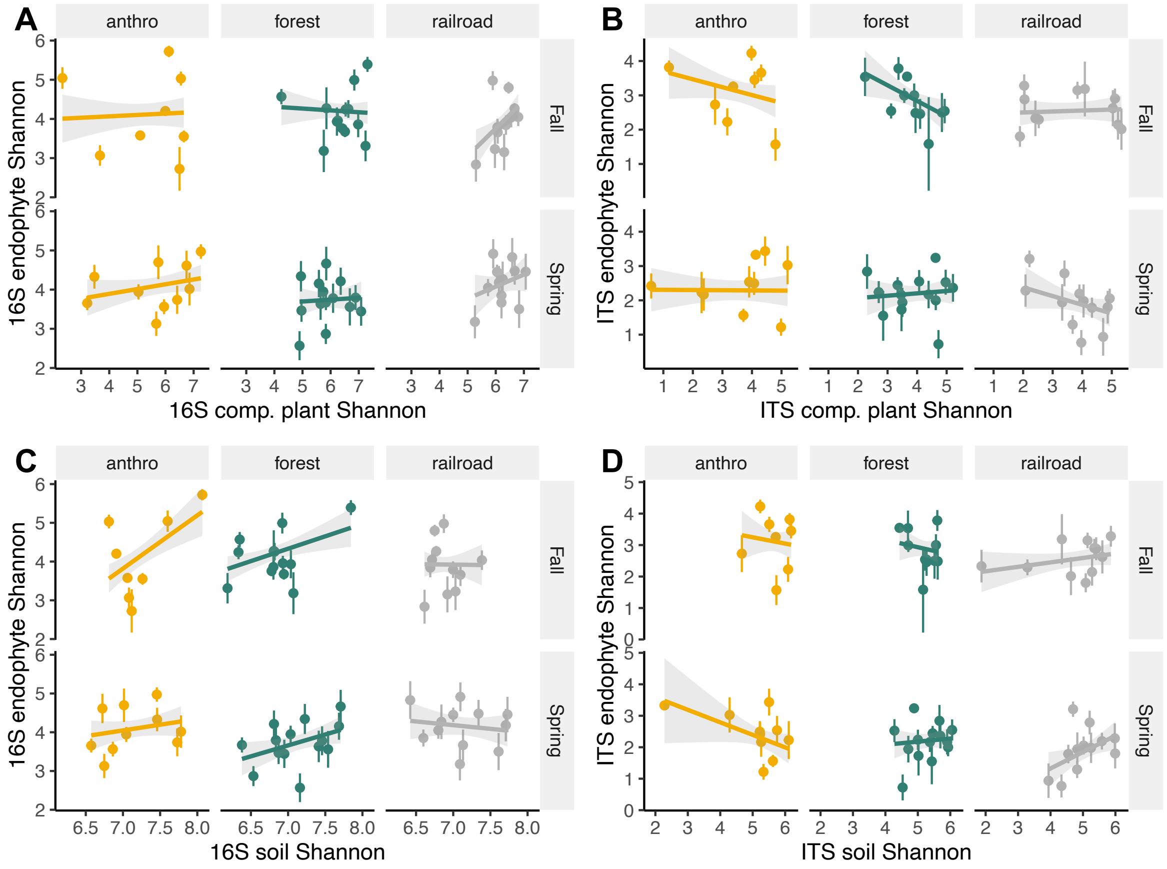


**Supp. Fig. R15:** Correlations between endophytes, companion plants, and soil for Shannon diversity in the microbiome. All plots have points colored by land use type, with points representing mean +/- standard error. Plots are faceted by land use and season. A) Correlations between companion plant and endophyte bacterial diversity. B) Correlations between companion plant and endophyte fungal diversity. C) Correlations between soil and endophyte bacterial diversity. D) Correlations between soil and endophyte fungal diversity.

**Supp. Table R19:** Fixed effects for the analysis of correlations between *A. thaliana* endophyte microbial Shannon diversity and microbial diversity in the environmental reservoirs of soil and companion plant microbiomes. Significance was evaluated using Type II or III Wald $X^2$ tests with Kenward-Roger degrees of freedom for two- or three-way interactions, respectively.

| *16S: Endophyte-Comp. plant bacterial diversity* | | | |
|---|---|---|---|
| | **Chisq** | **Df** | **Pr(>Chisq)** |
| Comp. plant bacterial Shannon diversity | 5.1507 | 1 | 0.02324* |
| Land use | 1.7505 | 2 | 0.41676 |
| Season | 2.0319 | 1 | 0.15403 |
| Plant size | 4.4248 | 1 | 0.03542* |
| Comp. plant diversity * Land use | 8.8377 | 2 | 0.01205* |
| ***ITS: Endophyte-Comp. plant fungal diversity*** | | | |
| | **Chisq** | **Df** | **Pr(>Chisq)** |
| (Intercept) | 37.5467 | 1 | 8.93E-10*** |
| Comp. plant fungal Shannon diversity | 2.0698 | 1 | 0.150237 |
| Land use | 5.5262 | 2 | 0.063096 |
| Season | 8.4088 | 1 | 0.003734** |
| Plant size | 4.4181 | 1 | 0.03556* |
| Comp. plant fungal diversity * Land use | 3.4382 | 2 | 0.17923 |
| Comp. plant fungal diversity * Season | 2.6213 | 1 | 0.105436 |
| Land use * season | 11.9112 | 2 | 0.002591** |
| Comp. plant fungal diversity * Land use * season | 11.9096 | 2 | 0.002593** |
| ***16S: Endophyte-soil bacterial diversity*** | | | |
| | **Chisq** | **Df** | **Pr(>Chisq)** |
| Soil bacterial Shannon diversity | 26.92 | 1 | 2.12E-07*** |
| Land use | 0.329 | 2 | 0.848322 |
| Season | 7.167 | 1 | 0.007425** |
| Plant size | 2.461 | 1 | 0.116701 |
| ***ITS: Endophyte-soil bacterial diversity*** | | | |
| | **Chisq** | **Df** | **Pr(>Chisq)** |
| (Intercept) | 0.4584 | 1 | 0.49836 |
| Soil fungal Shannon diversity | 0.4688 | 1 | 0.49353 |
| Land use | 0.1195 | 2 | 0.94198 |
| Season | 2.4419 | 1 | 0.11814 |
| Plant size | 2.8649 | 1 | 0.09053 |
| Soil fungal diversity * Land use | 0.2699 | 2 | 0.87374 |
| Soil fungal diversity * Season | 4.0237 | 1 | 0.04487* |
| Land use * Season | 6.3916 | 2 | 0.04093* |
| Soil fungal diversity * Land use * Season | 7.1804 | 2 | 0.02759* |

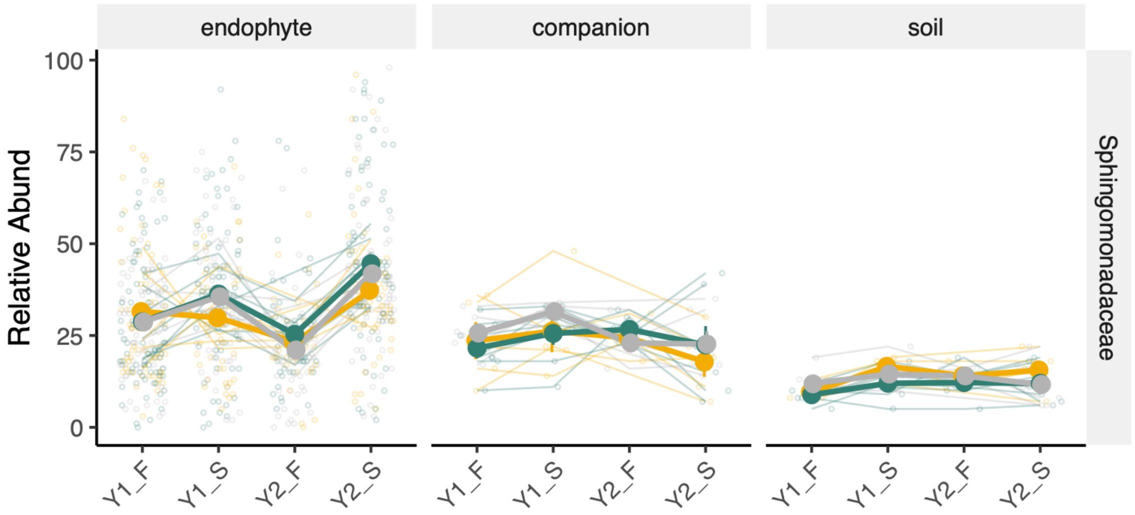


**Supp. Fig. R16:** Temporal change in Sphingomonadaceae relative abundance across the sampling seasons for the three different sample types. Points in endophytes represent individual *A. thaliana* endophytes, while companion plant and soil are pools per site and season. Thick lines and points +/- standard error show the average from each land use category.

**Supp. Table R20:** Fixed effects for analysis of microbial sharing on the most abundant bacterial family, Sphingomonadaceae (Sphingo.), between *A. thaliana* endophytes, companion plant, and soil microbiomes. Significance was evaluated using Type II or III Wald $X^2$ tests with Kenward-Roger degrees of freedom for two- or three-way interactions, respectively.

| ***Endophyte vs Companion plant*** | | | |
|---|---|---|---|
| | **Chisq** | **Df** | **Pr(>Chisq)** |
| (Intercept) | 70.8333 | 1 | < 2.20E-16*** |
| Comp. plant Sphingo | 15.0336 | 1 | 0.0001056*** |
| Land use | 10.3179 | 2 | 0.0057479** |
| Season | 10.7638 | 1 | 0.0010351** |
| Comp. plant Sphingo * Land use | 9.6138 | 2 | 0.008173** |
| Comp. plant Sphingo * Season | 13.5383 | 1 | 0.0002337*** |
| Land use * Season | 18.1234 | 2 | 0.000116*** |
| Comp. plant Sphingo * Land use * Season | 14.9701 | 2 | 0.0005614*** |
| ***Endophyte vs Soil*** | | | |
| | **Chisq** | **Df** | **Pr(>Chisq)** |
| Soil Sphingo | 2.9447 | 1 | 0.0861596 |
| Land use | 0.2914 | 2 | 0.8644332 |
| Season | 72.7163 | 1 | < 2.20E-16*** |
| Soil Sphingo * Land use | 15.7208 | 2 | 0.0003857*** |
| ***Companion plant vs Soil*** | | | |
| | **Chisq** | **Df** | **Pr(>Chisq)** |
| Soil Sphingo | 0.0928 | 1 | 0.7607 |
| Season | 0.0151 | 1 | 0.9021 |
| Land use | 1.6863 | 2 | 0.4304 |

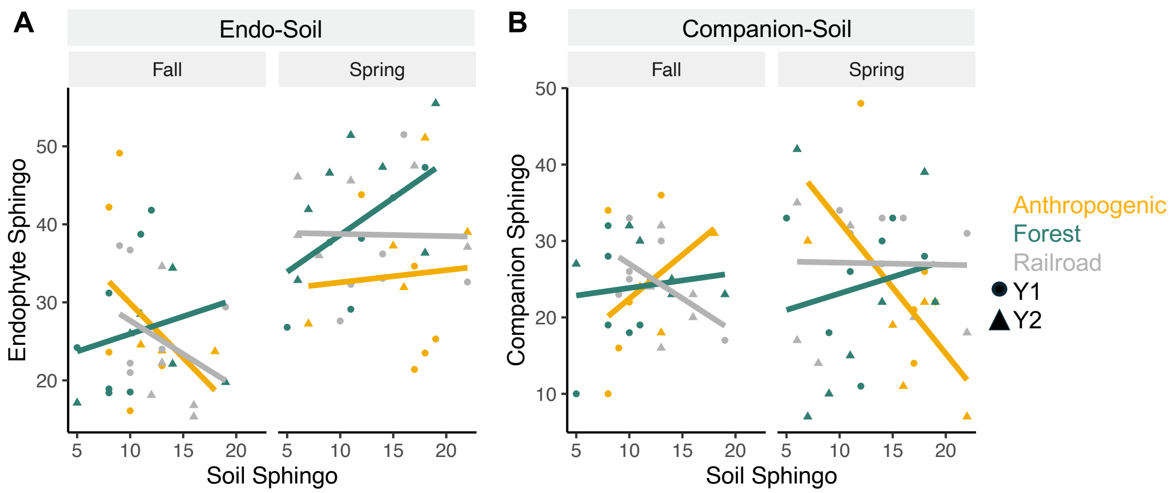


**Supp. Fig. R17:** Correlations for the relative abundance of the dominant bacterial family, Sphingomonadaceae, between A) soil and *A.* thaliana endophytes, and B) soil and companion plant microbiomes. For endophyte-soil comparison, the correlation between soil and mean +/- standard error for relative abundance in the endophytes for each population.

**Supp. Table R21A:** Fixed effects for analysis on the associations between disease symptoms PC1 and microbial diversity. Significance was evaluated using Type II Wald $X^2$ tests with Kenward-Roger degrees of freedom.

| | Chisq | Df | Pr(>Chisq) |
|---|---|---|---|
| Endophyte bacterial diversity | 8.8847 | 1 | 0.002876** |
| log10(endophyte bacterial load) | 4.0707 | 1 | 0.043634* |
| Land use | 0.5849 | 2 | 0.746446 |
| Comp. plant community diversity | 4.1929 | 1 | 0.040594* |
| Season | 19.0687 | 1 | 1.26E-05*** |
| Plant size | 7.2634 | 1 | 0.007038** |
| log10(bacterial load) * Land use | 5.1264 | 2 | 0.077057 |

**Supp. Table R21B:** Model selection for the association between disease symptoms PC1 and microbial diversity. The random effects (1|Year:Population) were the same across all models, but were removed in the table below to facilitate readability. The best model, M3, which matches the Table R21A is bolded.

M1: Disease PC1 ~ log10(bacterial load) * Land use + Season + Plant size

M2: Disease PC1 ~ Bacterial diversity + Land use + Season + Plant size

**M3: Disease PC1 ~ Bacterial diversity + log10(bacterial load) * Land use + comp. plant comm. diversity + Season + Plant size**

M4: Disease PC1 ~ log10(fungal load) + Land use + Season + Plant size

M5: Disease PC1 ~ Fungal diversity * Land use * Season + Plant size

M6: Disease PC1 ~ log10(fungal load) + Land use + Season + comp. plant comm. diversity + Plant size

| | npar | AIC | BIC | logLik | -2logLik | Chisq | Df | Pr(>Chisq) |
|---|---|---|---|---|---|---|---|---|
| M1 | 10 | 722.35 | 763.9 | -351.18 | 702.35 | 5.7075 | 1 | 0.0168923* |
| M2 | 8 | 716.85 | 750.09 | -350.42 | 700.85 | | | |
| **M3** | **12** | **711.41** | **761.27** | **-343.71** | **687.41** | **14.9388** | **2** | **0.0005703*** ** |
| M4 | 8 | 730.95 | 764.19 | -357.47 | 714.95 | 0 | 0 | |
| M5 | 15 | 720.31 | 782.63 | -345.15 | 690.31 | 0 | 3 | 1 |
| M6 | 9 | 726.06 | 763.45 | -354.03 | 708.06 | 6.8867 | 1 | 0.0086838** |

**Supp. Table R22A:** Fixed effects for analysis on the associations between disease symptoms PC2 and microbial diversity. Significance was evaluated using Type II Wald $X^2$ tests with Kenward-Roger degrees of freedom.

| | **Chisq** | **Df** | **Pr(>Chisq)** |
|---|---|---|---|
| log10(fungal load) | 5.4184 | 1 | 0.019925* |
| Season | 79.7005 | 1 | < 2.20E-16*** |
| Land use | 9.3463 | 2 | 0.009343** |
| Plant size | 2.5865 | 1 | 0.107776 |
| log10(fungal load) * Season | 9.8157 | 1 | 0.00173** |

**Supp. Table R22B:** Model selection for the association between disease symptoms PC1 and microbial diversity. The random effects (1|Year:Population) were the same across all models, but were removed in the table below to facilitate readability. The best model, M4, which matches the Table R22A is bolded.

M1: Disease PC2 ~ log10(bacterial load) + Land use + Season + Plant size

M2: Disease PC2 ~ bacterial diversity + Land use * Season + Plant size

M3: Disease PC2 ~ bacterial diversity + log10(bacterial load) * Land use + comp. plant comm. diversity + Season + Plant size

**M4: Disease PC2 ~ log10(fungal load) * Season + Land use + Plant size**

M5: Disease PC2 ~ fungal diversity + Land use + Season + Plant size

M6: Disease PC2 ~ log10(fungal load) * Season Land use + comp. plant comm. diversity + Plant size

| | **npar** | **AIC** | **BIC** | **logLik** | **-2logLik** | **Chisq** | **Df** | **Pr(>Chisq)** |
|---|---|---|---|---|---|---|---|---|
| M1 | 8 | 481.48 | 514.71 | -232.74 | 465.48 | | | |
| M2 | 10 | 476.58 | 518.13 | -228.29 | 456.58 | 0 | 1 | 1 |
| M3 | 12 | 481.49 | 531.35 | -228.75 | 457.49 | 0 | 2 | 1 |
| **M4** | **9** | **468.68** | **506.08** | **-225.34** | **450.68** | **13.0297** | **1** | **0.0003066*** ** |
| M5 | 8 | 479.71 | 512.95 | -231.86 | 463.71 | 1.7632 | 0 | |
| M6 | 10 | 469.45 | 511 | -224.72 | 449.45 | 7.128 | 0 | |

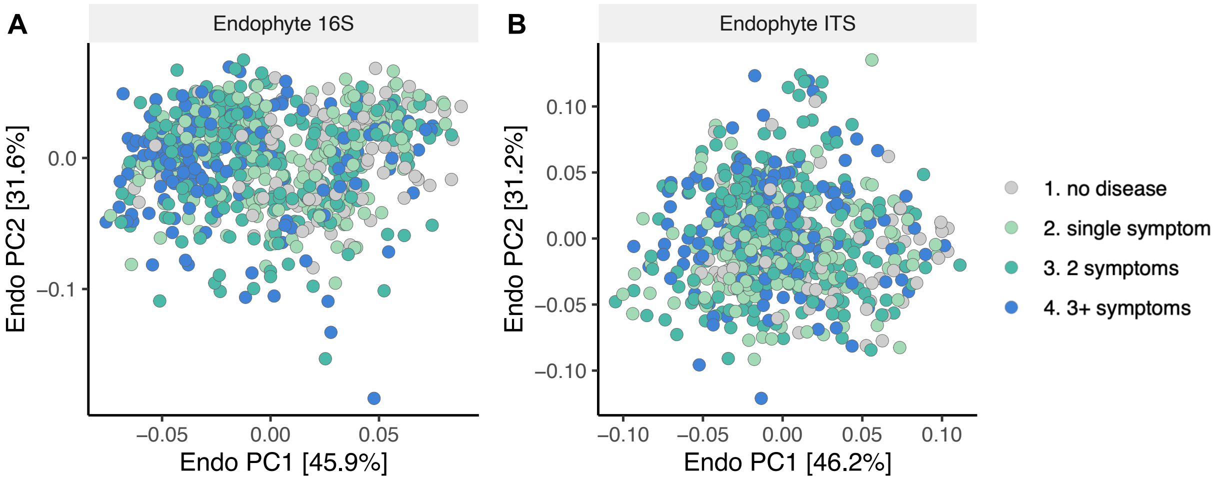


**Supp. Fig. D1:** Disease does not structure the *A. thaliana* endophytic microbiome. A) In the context of season and land use, disease PC1 explained significant but very low amounts of variance in bacterial microbiomes ($R^2 = 0.007$, $F_{1,743} = 11.43$, $p = 0.001$). B) Disease did not explain any significant variance for fungal microbiome structure ($F_{1,624} = 2.36$, $p = 0.11$).

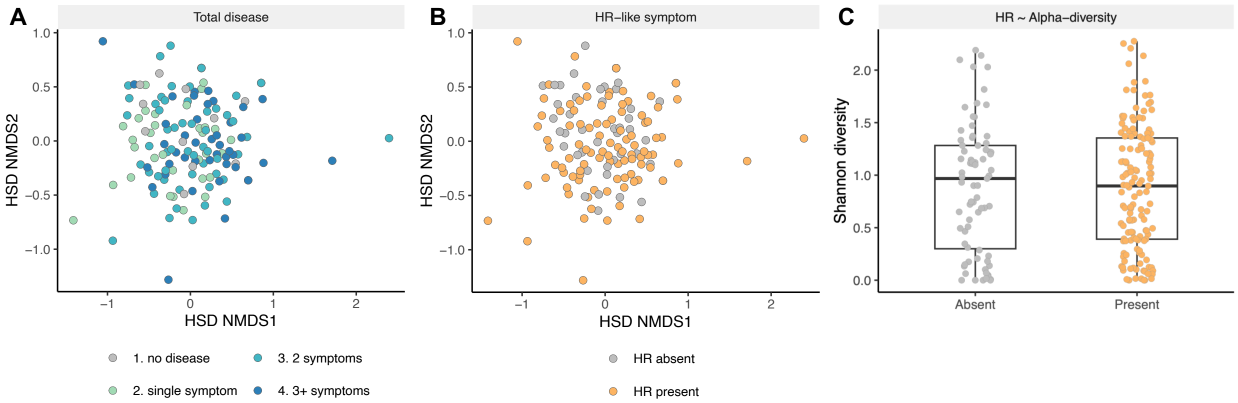


**Supp. Fig. D2:** No clear relationship between disease and *Pseudomonas* community structure. A) Disease does not structure *Pseudomonas* beta-diversity (PERMANOVA, $F_{1,140}$ = 1.69, p = 0.149), suggesting that *HSD* amplicon did not capture pathogens associated with disease severity. Points are shown in NMDS based on weighted UniFrac distance, colored by the number of disease symptoms, which corresponds to disease PC1. B) There is also no difference between plants with or without HR-like symptoms (PERMANOVA, $F_{1,140}$ = 0.913, p = 0.454). C), and no difference in *HSD* Shannon diversity between plants with and without HR-like symptoms (mixed linear model, beta = -0.009 +/- 0.08 SE, t = -0.11, p = 0.916).

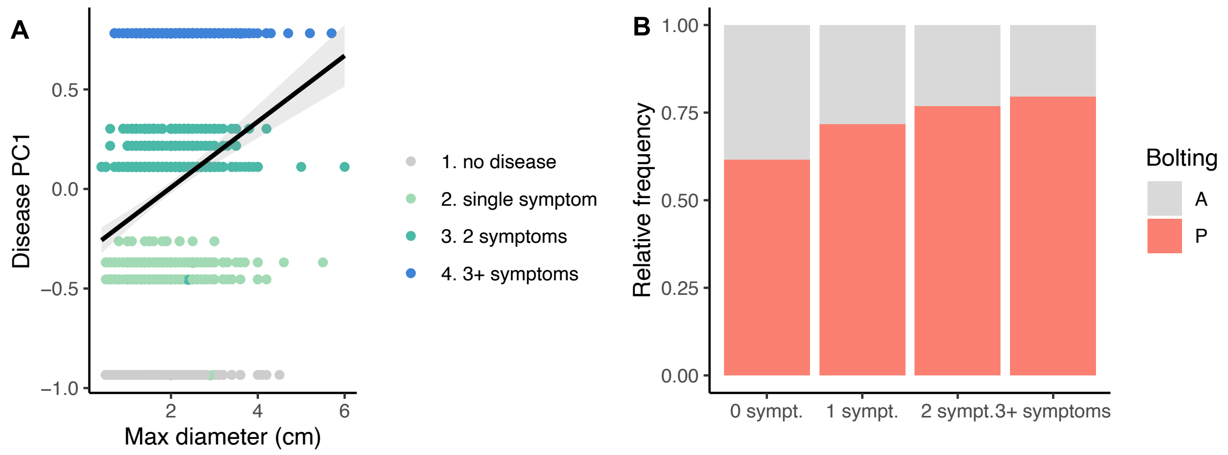


**Supp. Fig. D3:** Diseased plants do not pay an obvious cost on infection. A) Larger plants have more disease symptoms, as disease PC1 represents the number of symptoms. Points represent individual plants, colored by the number of symptoms observed. B) The number of symptoms does not impact the frequency of plants that are bolting. A = no bolting, P = bolting. The effects were compared only in the spring, as bolting does not occur in the fall.

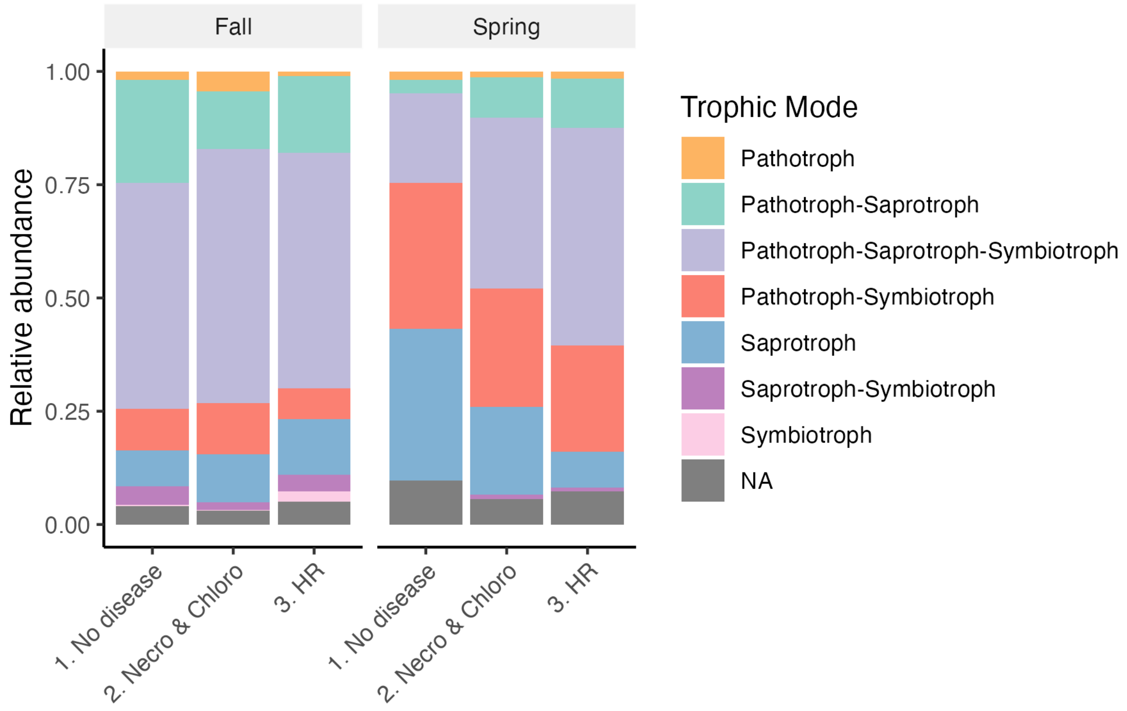


**Supp. Fig. D4**: Pathotrophs do not vary consistently across disease symptom groups or season. There is a marginal increase in pathotrophs (orange) in the "Necro & Chloro" groups, but the largest change is associated with an increase in saprotrophs (blue) in the "No disease" plants in the spring.

***REFERENCES***

9. Rice P, Bleasby A, Ison J. In press. *The EMBOSS Users Guide*. Cambridge University Press.

10. Kõressaar T, Lepamets M, Kaplinski L, Raime K, Andreson R, Remm M. 2018 Primer3_masker: integrating masking of template sequence with primer design software. *Bioinformatics* **34**, 1937–1938.

11. Broad Institute. 2019 *Picard Toolkit*. See https://broadinstitute.github.io/picard/.

12. Leach R, Parsons L. 2019 Barcode Splitter, version 0.18.5 [Software].
